# Ascorbate is a physiological tumor suppressor metabolite in leukemia

**DOI:** 10.64898/2026.10.20.760526

**Authors:** Tobias Wijshake, Joseph Rose, Ji Hyung Jun, Edward Kwarteng, Xiaotong Yang, Jacob Zielke, Swetha Mahesula, Li Li, Stephen S. Chung, Stefano Comazzetto, Sean J. Morrison, Weina Chen, Michalis Agathocleous

## Abstract

Accumulation of oncometabolites promotes tumorigenesis, but it is not known if there are naturally occurring tumor suppressor metabolites whose deficiency within cancer-initiating cells causes cancer. Many leukemia patients are ascorbate deficient. Here we show that hematopoietic-specific ascorbate depletion in mice is sufficient to cause or promote leukemogenesis. Hematopoietic-specific ascorbate depletion promoted leukemogenesis by acting within leukemia initiating cells. Ascorbate depletion in myeloid leukemia expanded plasmacytoid dendritic cells, an effect mediated by the ascorbate molecular targets TET2 and TET3. Leukemia cells accumulated ascorbate as part of a tumor suppressor response to neoplastic proliferation that limited progression but depleted circulating ascorbate. Ascorbate restoration in mice with established leukemias extended survival. Hematopoietic-specific ascorbate depletion caused leukemia in collaboration with *TET2* loss and was sufficient to cause leukemia with high penetrance even in the absence of other engineered mutations. These results identify ascorbate as a physiological tumor suppressor metabolite in blood cancers.

## INTRODUCTION

Variations in metabolite levels regulate cancer development. Dietary depletion of several amino acids^1^, or experimental elevation of ascorbate, α-ketoglutarate, or β-hydroxybutyrate^2-5^ can inhibit cancer growth. A few endogenous metabolites, termed oncometabolites, are naturally elevated in cancer-initiating cells and collaborate with cancer-driver mutations to promote tumorigenesis^6^. Known oncometabolites transform cells by inhibiting α-ketoglutarate dependent dioxygenases (αKGDDs) that modify nucleic acids or histones. It remains unknown whether there are naturally occurring tumor suppressor metabolites that at physiological levels suppress transformation *in vivo*. If such metabolites exist, then their depletion from cancer-initiating cells should collaborate with cancer-driver mutations to cause cancer.

Ascorbate is a naturally occurring αKGDD activator^7^. Its supraphysiological elevation by exogenous administration inhibits the growth of many cancers in mice^5,8^ by causing oxidative stress^5,9^, damaging DNA^10-13^, or activating αKGDDs^14-16^. Blood cancer cells are particularly sensitive to supraphysiological ascorbate^10,15,17-22^. These findings establish that supraphysiological ascorbate has potent anti-cancer activity but leave open the question of whether ascorbate is a tumor-suppressor metabolite at physiological levels. Ascorbate inhibits the transformation of nonmalignant fibroblasts^23^ and the self-renewal of pre-leukemic stem cells^15^ in culture, raising the possibility that it limits cancer development *in vivo*.

Little is known about the physiological role of naturally occurring ascorbate in cancer because few studies have depleted it in cancer models. Systemic ascorbate depletion in *Gulo^−/−^* mice reduces activity of TET2, a leukemia tumor suppressor, and accelerates myeloid leukemia development^24^. Ascorbate has not been depleted in a cell-type specific manner in leukemia or other cancer models^8,24^. Therefore, it is not known if ascorbate acts cell-autonomously within leukemia-initiating cells or by modulating niche cells or other cell types.

Whether ascorbate naturally modulates cancer development is relevant to human cancer because many humans have low plasma ascorbate levels. Most mammals synthesize ascorbate in the liver via L-Gulonolactone oxidase (GULO), but humans are GULO-deficient, an evolutionary gene loss that turned ascorbate into an essential dietary micronutrient better known as vitamin C^25,26^. Consequently, plasma ascorbate concentrations vary 10-fold in western countries^27,28^ and people with levels in the lowest quartile have higher cancer mortality^28,29^. Ascorbate depletion is highly prevalent in patients with blood neoplasms as compared to the general population^17,30-36^. Ascorbate supplementation of patients with early-stage myeloid neoplasms or clonal cytopenias who had below-average ascorbate levels at baseline may promote survival^37^.

### Slc23a2 is the physiological ascorbate transporter in leukemia cells

Ascorbate synthesized in the liver or taken up from the diet enters cells via its dedicated transporters SLC23A1/A2^38,39^ (**Fig. 1A**). SLC23A1 is expressed mainly in tissues involved in bulk transport, and SLC23A2 in most tissues^38^. When oxidized, ascorbate can enter cells via GLUT transporters, a mechanism used by some cancer cells *in vitro*^40^. It is not known if *in vivo* ascorbate is imported in cancer cells via SLC23A1/A2 or GLUT transporters. *Slc23a2* is required in mouse hematopoietic stem and progenitor cells (HSPCs) for ascorbate uptake^24,41^.

**Figure 1.**
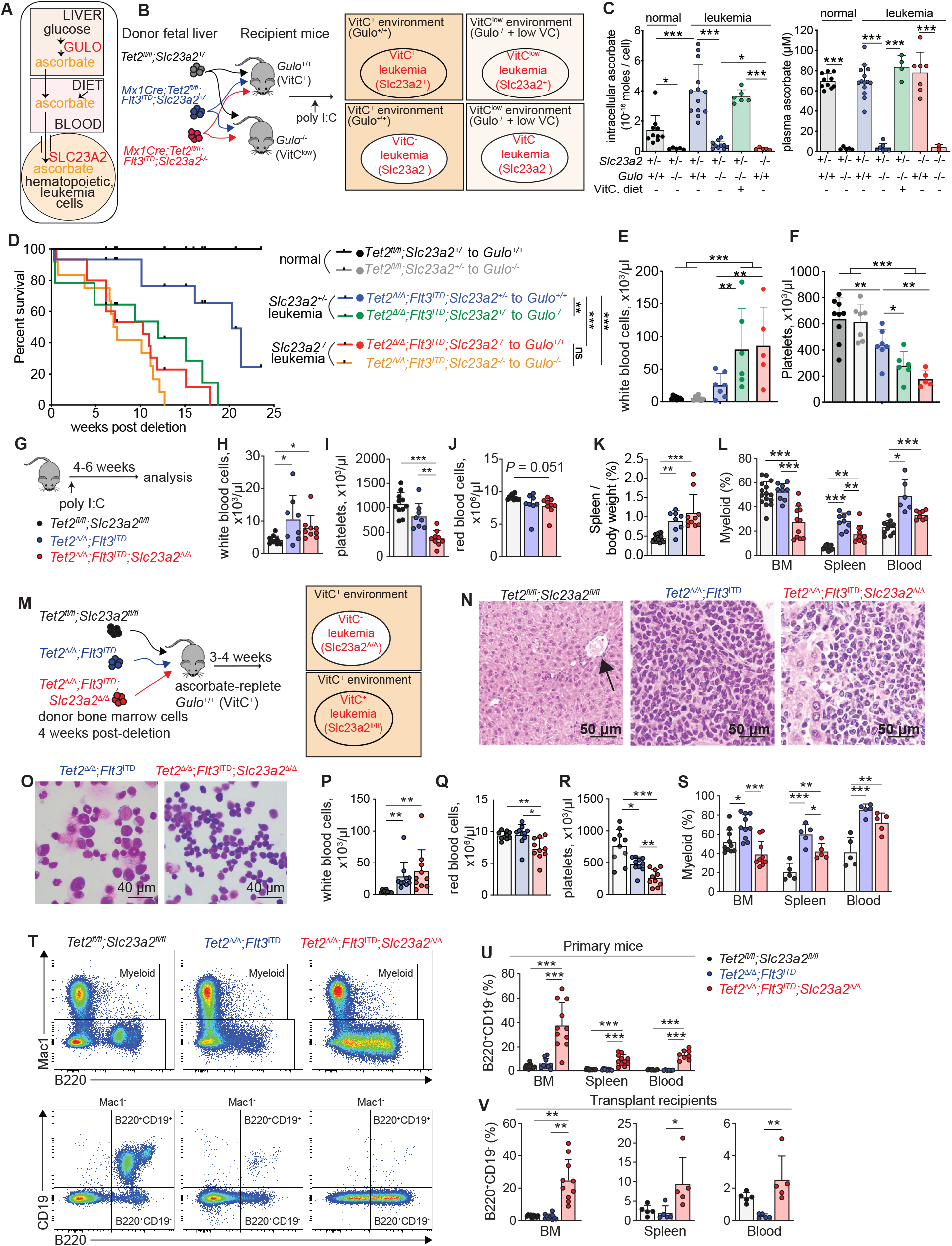
Cell autonomous ascorbate depletion promotes leukemogenesis. (**A**) Ascorbate is synthesized in the liver or consumed in the diet, circulates, and enters cells via SLC23A2. (**B-F**) Transplantation of ascorbate-replete or ascorbate-depleted leukemia cells into ascorbate-replete or ascorbate-depleted mice to assess cell-autonomous vs non-cell-autonomous role of ascorbate in leukemogenesis. (**C**) Ascorbate levels in the plasma or intracellularly in normal or leukemic cells (*Tet2^Δ/Δ^;Flt3^ITD^*or *Tet2^Δ/+^;Flt3^ITD^*) in transplant recipients. (**D-F**) Kaplan-Meier survival curve of transplant recipient mice and blood cell counts from the indicated genotypes (survival curve, n=12-19 mice/genotype). (**G-L**) Analysis of mice of the indicated genotypes for blood cell counts, spleen size, and myeloid cell frequencies. (**M-T**) Transplantation of ascorbate-replete or ascorbate-depleted leukemia cells into ascorbate-replete wild-type mice and analysis 4-8 weeks later. (**N**) Images of liver H&E staining. Normal liver is composed of cords of hepatocytes with abundant cytoplasm and portal triads (black arrow – left). Liver in leukemic mice is infiltrated by sheets of medium-sized to large cells with irregular nuclei, variable nucleoli, and scant to moderate cytoplasm (middle and right). (**O**) Images of Giemsa-stained bone marrow smears. *Tet2^Δ/Δ^*;*Flt3^ITD^* smears present scattered medium-sized to large cells (20-30% of total) with moderately dispersed chromatin and scant to moderate cytoplasm with some cytoplasmic granules on a background of normal cells. *Tet2^Δ/Δ^*;*Flt3^ITD^*;*Slc23a2^Δ/Δ^*smears present a predominant population of lymphoid-appearing medium-sized cells (60-70% of total) with slightly irregular nuclei, variable or inconspicuous nucleoli, and scant cytoplasm. (**P-S**) Blood cell counts and analysis of mice from the indicated genotypes. (**T**) Representative flow plots showing elevated myeloid and B220^+^CD19^−^ cells in *Tet2^Δ/Δ^;Flt3^ITD^;Slc23a2^Δ/Δ^* bone marrow. (**U-V**) Frequencies of B220^+^CD19^−^ cells in primary and transplant recipient mice. In all figures, data represent mean ± s.d., \**p* < 0.05, \*\**p* < 0.01, \*\*\**p* < 0.001 and individual dots in bar graphs in all figures represent biological replicates (mice). Statistical significance was assessed with a Brown-Forsythe ANOVA (C, L, spleen and blood; U, and V, BM), Mantel-Cox test (D), one-way ANOVA (E-F, I-J, L, BM; R-S, and V, blood), or a Kruskal-Wallis test (H, K, P-Q and V, spleen). Statistical tests for all figures were selected as described in the methods section by testing if the data from different treatments were consistent with assumptions of normality or equality of variance.

We crossed *Slc23a2^+/−^* mice^39^ with *Mx1Cre;Tet2^fl/fl^;Flt3^ITD^*mice which develop acute myeloid leukemia (AML)^42^. We transplanted *Mx1Cre;Tet2^fl/fl^;Flt3^ITD^* fetal liver cells that were *Slc23a2^−/−^* or *Slc23a2^+/−^*to irradiated wild-type or *Gulo^−/−^* recipients to compare the effects of cell-autonomous or systemic ascorbate depletion on leukemogenesis (**Fig. 1B**)*. Slc23a2^−/−^*leukemia cells growing in wild-type mice had no detectable intracellular ascorbate (**Fig. 1C**), suggesting SLC23A2 is the physiological ascorbate transporter in leukemia cells.

We provided *Gulo^−/−^* mice with low dietary ascorbate to induce systemic ascorbate depletion but avoid scurvy^24^. AML cells growing in *Gulo^−/−^* mice had low intracellular ascorbate that was restored to normal levels by 1% ascorbate chow (**Fig. 1C**). This indicated that ascorbate levels in leukemia cells respond to changes in dietary and systemic ascorbate levels. AML cells had higher intracellular ascorbate levels compared to normal hematopoietic cells (**Fig. 1C**). These results suggest that leukemia cells import circulating ascorbate via SLC23A2 and accumulate it compared to normal cells.

### Cell-autonomous ascorbate depletion promotes leukemogenesis

*Slc23a2* deletion enhances the function of HSCs and multipotent progenitors (MPP)^24,41^ but its effect on leukemogenesis is unknown. We compared leukemogenesis in the orthogonal models of systemic or cell-autonomous ascorbate depletion (**Fig. 1A-B**). *Gulo^−/−^* recipients of *Tet2^Δ/Δ^;Flt3^ITD^*leukemia cells succumbed to leukemia faster than wild-type recipients (**Fig. 1D**), consistent with our previous data^24^. Wild-type recipients of *Tet2^Δ/Δ^;Flt3^ITD^;Slc23a2^−/−^*leukemia succumbed faster than wild-type recipients of *Tet2^Δ/Δ^;Flt3^ITD^;Slc23a2^+/−^* cells. The magnitude of the effect of *Slc23a2* deletion was similar to that of *Gulo* deletion (**Fig. 1D**). Therefore, ascorbate depletion accelerates leukemia development, and this is mainly or entirely due to action of ascorbate within hematopoietic or leukemia cells. *Slc23a2^−/−^;Tet2^Δ/Δ^;Flt3^ITD^* leukemia in wild-type recipients, or *Tet2^Δ/Δ^;Flt3^ITD^* leukemia in *Gulo^−/−^* recipients, caused increased leukocytosis, thrombocytopenia, and anemia compared to ascorbate-replete *Tet2^Δ/Δ^;Flt3^ITD^*leukemia, consistent with accelerated leukemogenesis (**Fig. 1E-F, Fig. S1A-B**). Ascorbate-replete or depleted *Tet2^Δ/Δ^;Flt3^ITD^* leukemias were characterized by myeloid expansion in the blood, liver, and spleen (**Fig. S1C).**

We generated *Mx1Cre;Slc23a2^fl/fl^;Tet2^fl/fl^;Flt3^ITD^*mice to test the effects of adult *Slc23a2* deletion in leukemia. Mice were treated with poly I:C to delete *Tet2* and *Slc23a2* and analyzed 4-6 weeks later (**Fig. 1G**). Similar to *Tet2^Δ/Δ^;Flt3^ITD^* mice, *Slc23a2^Δ/Δ^;Tet2^Δ/Δ^;Flt3^ITD^*mice developed leukemia marked by leukocytosis, thrombocytopenia, anemia, splenomegaly and B cell depletion (**Fig. 1H-K, Fig. S1D-G**). The myelomonocytic expansion that characterizes *Tet2^Δ/Δ^;Flt3^ITD^* mice^24,42^ was present but blunted in *Slc23a2^Δ/Δ^;Tet2^Δ/Δ^;Flt3^ITD^* mice (**Fig. 1L, Fig. S1H**). To avoid *Mx1Cre-*mediated deletion in other tissues, we assessed leukemogenesis by transplanting *Slc23a2^Δ/Δ^;Tet2^Δ/Δ^;Flt3^ITD^*or *Tet2^Δ/Δ^;Flt3^ITD^* hematopoietic cells into irradiated wild-type mice (**Fig. 1M**). Recipient mice developed leukemia infiltrating the liver and bone marrow (**Fig. 1N-O**). *Slc23a2* deletion exacerbated thrombocytopenia and anemia, consistent with leukemia acceleration, but blunted the myeloid expansion seen in *Tet2^Δ/Δ^;Flt3^ITD^* leukemias (**Fig. 1P-T**). *Slc23a2^Δ/Δ^;Tet2^Δ/Δ^;Flt3^ITD^*primary or transplant recipient mice showed a distinct leukemic cell morphology (**Fig. 1O**) and expansion of a B220^+^CD19^−^ cell population (**Fig. 1T-V, Fig. S1I-K**).

### Ascorbate depletion promotes the generation of a plasmacytoid dendritic cell neoplasm associated with AML

B220^+^CD19^−^ cells include plasmacytoid dendritic cells (pDCs). B220^+^CD19^−^SiglecH^+^PDCA^+^, B220^+^CD19^−^SiglecH^−^PDCA^+^ and B220^−^CD19^−^SiglecH^+^PDCA^+^ pDCs expanded in the bone marrow, spleen, and blood, of *Slc23a2^Δ/Δ^;Tet2^Δ/Δ^;Flt3^ITD^*leukemias, in both primary mice and transplant recipients (**Fig. 2A-C, Fig. S2A-E**). Expanded pDCs characterize several blood neoplasms, including a subtype of AML (pDC-AML), mature pDC proliferations associated with myeloid neoplasms (MPDMN), and blastic pDC neoplasms (BPDCN)^43^. Immunofluorescence analysis showed abundant B220^+^SiglecH^+^PDCA^+^ cells scattered through the *Slc23a2^Δ/Δ^;Tet2^Δ/Δ^;Flt3^ITD^* bone marrow, interspersed with other cells (**Fig. 2D**). The combination of pDC and monocytic expansion and the interspersed appearance of marrow pDCs in *Slc23a2^Δ/Δ^;Tet2^Δ/Δ^;Flt3^ITD^*mice were more consistent with pDC-AML rather than MPDMN, in which pDCs typically form nodules^44^.

**Figure 2.**
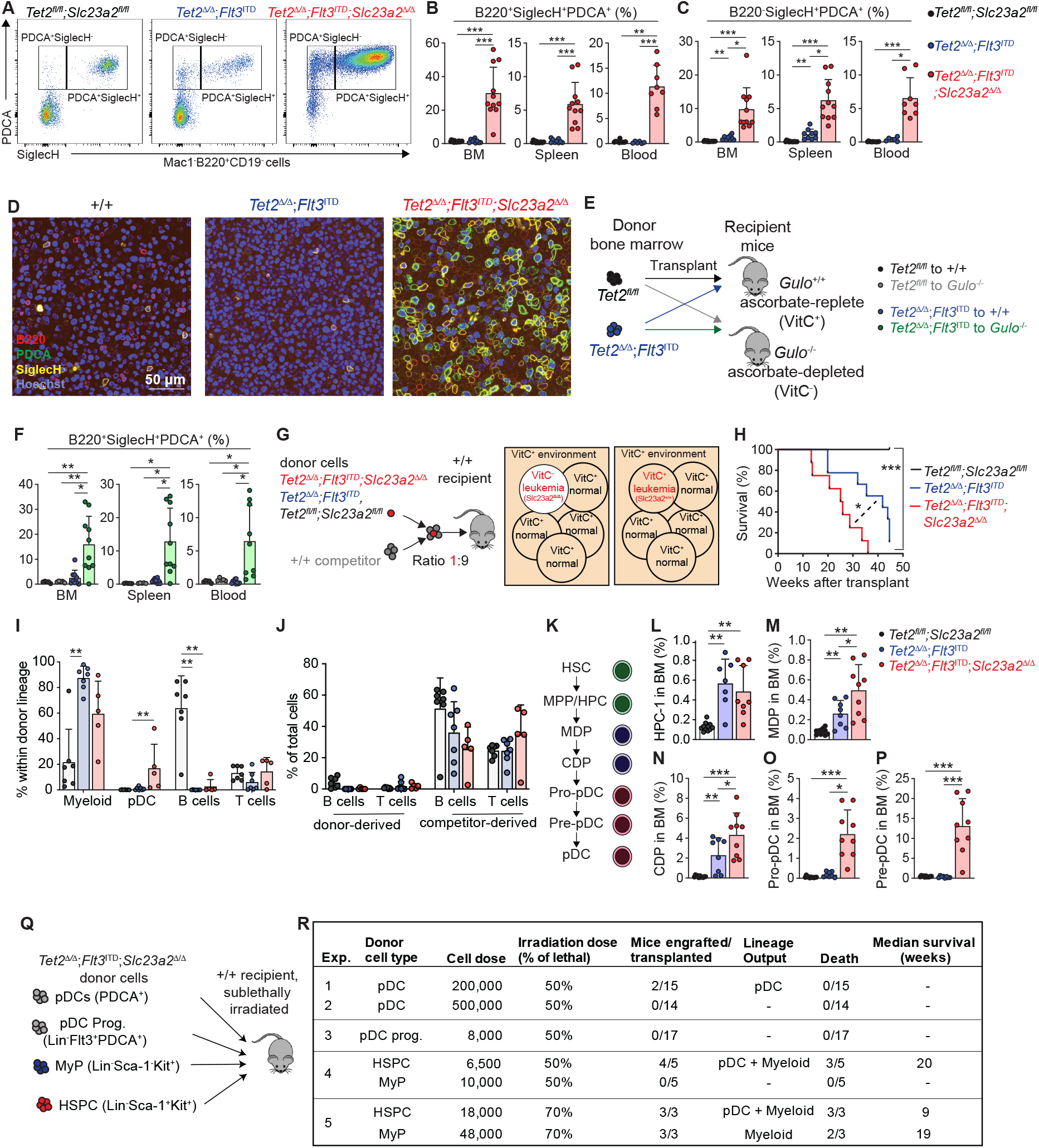
Ascorbate depletion promotes the generation of a plasmacytoid dendritic cell neoplasm. (**A-C**) Representative flow plots from the bone marrow and frequencies of pDC populations in primary mice. (**D**) Representative immunofluorescence images of whole-mount bone marrow stained with antibodies against B220, PDCA, and SiglecH. (**E-F**) Transplantation of ascorbate-replete leukemia cells into ascorbate-depleted mice followed by analysis 5 weeks later. (**G-J**) 40,000 bone marrow cells from the indicated genotype were mixed with 360,000 wild-type competitor cells and transplanted into lethally irradiated wild-type mice. Mice with > 1% reconstitution were included in the analysis. (**H**) Kaplan-Meier survival curve of recipient mice (n = 8-10 mice/genotype). (**I**) Frequency within the donor lineage of the indicated cell populations in recipient blood 12 weeks after transplant. (**J**) Percentage of donor– or competitor-derived B and T cells in blood of recipient mice 12 weeks after transplant. (**K**) Schematic of pDC differentiation. (**L-P**) Frequencies of oligopotent and pDC lineage progenitors in the bone marrow. (**Q-R**) Transplants of purified cell populations to sublethally irradiated recipients to assess leukemia-initiating capacity and results. Statistical significance was assessed with a Brown-Forsythe ANOVA (B, BM and spleen; F, L-N, and P), Kruskal-Wallis test (B, blood; C, I, and O), or a Mantel-Cox test (H).

To test if the effects of *Slc23a2* deletion were recapitulated by an orthogonal method of ascorbate depletion, *Gulo^−/−^*mice completely deprived of dietary ascorbate were transplanted with *Tet2^Δ/Δ^;Flt3^ITD^*cells and analyzed 5 weeks later (**Fig. 2E**). Ascorbate depletion in *Gulo^−/−^* mice phenocopied *Slc23a2* deletion in promoting pDC and monocyte expansion (**Fig. 2F, Fig. S2F-J**). Therefore, SLC23A2 suppresses pDC-AML by importing ascorbate rather than an unrelated mechanism.

### The developmental origin of ascorbate-depleted AML

To test if ascorbate suppresses leukemogenesis by acting cell-intrinsically in leukemia-initiating cells or non-cell intrinsically in other cells, such as a normal hematopoietic or immune cell, we transplanted *Slc23a2^Δ/Δ^;Tet2^Δ/Δ^;Flt3^ITD^*or *Tet2^Δ/Δ^;Flt3^ITD^* bone marrow cells with wild-type competitor hematopoietic cells at a ratio of 1:9 into irradiated recipients (**Fig. 2G**). This models the growth of leukemia from a minority clone in the presence of a wild-type hematopoietic system. Mice with *Slc23a2^Δ/Δ^;Tet2^Δ/Δ^;Flt3^ITD^*leukemia cells died earlier than mice with *Tet2^Δ/Δ^;Flt3^ITD^*leukemia cells, consistent with accelerated leukemogenesis (**Fig. 2H**). *Slc23a2^Δ/Δ^;Tet2^Δ/Δ^;Flt3^ITD^* leukemias had myeloid and pDC expansion, in contrast to *Tet2^Δ/Δ^;Flt3^ITD^*leukemias, which had a purely myeloid expansion (**Fig. 2I**). B and T cells originated from wild-type competitors (**Fig. 2J**). This indicated that ascorbate depletion promoted leukemogenesis in the presence of a normal ascorbate-replete adaptive immune system. Therefore, ascorbate depletion promotes leukemogenesis by acting cell-autonomously in leukemia-initiating cells rather than in other cell types.

pDC-AML is a pathologically distinct AML subtype characterized by pDC expansions, makes up ∼5% of total AML, and has adverse prognosis^43,45,46^. *TET2* or *FLT3* mutations are found in 15-25% of pDC-AML^45^. To understand the developmental origins of ascorbate-depleted pDC-AML, we analyzed the hematopoietic stem and progenitor compartment (**Fig. 2K**). At 4 weeks after deletion, *Slc23a2^Δ/Δ^;Tet2^Δ/Δ^;Flt3^ITD^*mice had depleted phenotypic CD150^+^CD48^−^Lin^−^Sca-1^+^Kit^+^ HSCs and CD150^−^CD48^−^Lin^−^Sca-1^+^Kit^+^ MPPs and expanded CD150^−^CD48^+^Lin^−^Sca-1^+^Kit^+^ oligopotent progenitors (HPC-1), similar to *Tet2^Δ/Δ^;Flt3^ITD^*mice^42^ (**Fig. 2L, Fig. S3A**). *Slc23a2* deficiency reduced common myeloid progenitors (CMPs) and granulocyte-monocyte progenitors (GMPs) suggesting a shift away from pure myeloid expansion (**Fig. S3B-E**). pDCs arise partly from the myeloid lineage via macrophage-dendritic cell progenitors (MDPs)^47^. We hypothesized that ascorbate depletion promoted pDC-AML by amplifying MDP and committed pDC progenitors when combined with *Tet2* deletion and *Flt3^ITD^*. *Slc23a2* deletion increased the frequencies of Lin^−^Kit^+^Sca-1^−^Flt3^+^CD115^+^SiglecH^−^ MDPs, Lin^−^Kit^−^CD115^+^Flt3^+^SiglecH^−^ common dendritic cell progenitors (CDPs), Lin^−^Kit^−^Flt3^+^CD115^+^SiglecH^+^ pro-pDCs and Lin^−^Kit^−^ Flt3^+^CD115^−^SiglecH^+^ pre-pDCs (**Fig. 2M-P, Fig. S3F**). Therefore, *Slc23a2* deletion amplifies pDCs in AML by expanding the earliest pDC progenitor cells.

*Slc23a2^Δ/Δ^;Tet2^Δ/Δ^;Flt3^ITD^* leukemia cells transplanted to wild type recipients, or *Tet2^Δ/Δ^;Flt3^ITD^*leukemia cells transplanted to *Gulo^−/−^* recipients completely deprived of ascorbate also expanded MDP and committed pDC progenitors (**Fig. S3G-Q**). The fact that the effects of *Slc23a2* deletion on leukemia acceleration (**Fig. 1D**, **Fig. 2H**) and pDC-AML development (**Fig. 2A-D**, **Fig. 2K-P**) were phenocopied by *Gulo* deletion indicated they were due to ascorbate depletion rather than an unknown SLC23A2 substrate.

Self-renewing leukemia stem cells (LSCs) sustain AML and cause relapse. They are experimentally defined by their ability to initiate disease in transplantation experiments. In the *Tet2^Δ/Δ^;Flt3^ITD^*AML model, LSCs are within Lin^−^Sca-1^+^Kit^+^ multipotent/oligopotent progenitors^42^. We tested which *Slc23a2^Δ/Δ^;Tet2^Δ/Δ^;Flt3^ITD^*cell fractions could transplant AML to recipient mice (**Fig. 2Q**). Purified *Slc23a2^Δ/Δ^;Tet2^Δ/Δ^;Flt3^ITD^*pDC-AML cell fractions sorted from primary recipients were transplanted to sublethally irradiated secondary recipients to test LSC activity. Purified PDCA^+^ pDC lineage cells produced pDCs in only 2/29 recipients and did not generate any leukemias (**Fig. 2R**). This is consistent with a prior report that pDCs in human pDC-AML cannot transplant disease and are instead produced by leukemic blasts^46^. Lin^−^Sca-1^−^ Kit^+^ restricted myeloid progenitors had some leukemia-initiating ability and produced myeloid cells but not pDCs (**Fig. 2R**). Purified Lin^−^Sca-1^+^Kit^+^ HSPCs had much higher leukemia initiating ability and generated lethal leukemias marked by progressive leukocytosis, anemia and myeloid/pDC expansion. These grew in the presence of recipient-derived, ascorbate-replete normal myeloid, B, and T cells (**Fig. 2R, Fig. S3R-T**). Therefore, LSC activity in pDC-AML is found within Lin^−^Sca-1^+^Kit^+^ HSPCs, and ascorbate depletion within LSCs in the presence of a normal hematopoietic system is sufficient to drive pDC-AML.

### Ascorbate suppresses leukemia development by activating Tet2 and Tet3

Ascorbate promotes the activity of α-ketoglutarate dependent dioxygenases, including Ten-Eleven Translocation 1-3 (TET1-3) nucleic acid methylcytosine dioxygenases. Inactivating *TET2* mutations drive the development of myeloid leukemia and other blood cancers^48^. Systemic ascorbate depletion in *Gulo^−/−^* mice enhances HSC function primarily by inhibiting TET2^24^ but in myeloid progenitors and leukemia cells it reduces TET activity and enhances leukemogenesis even in *Tet2-*null cells, suggesting it may inhibit multiple TET enzymes^24^. We investigated the molecular mechanisms by which hematopoietic ascorbate depletion promotes leukemogenesis.

To understand if ascorbate depletion cell-intrinsically reduces TET activity, we quantified levels of the TET product 5-hydroxymethylcytosine (5hmC) on the DNA of leukemia cells. *Tet2^Δ/Δ^;Flt3^ITD^* cells had reduced 5hmC in Lin^−^Sca-1^+^Kit^+^ HSPCs, GMPs, pDCs, and total CD45^+^ marrow cells as compared to wild type cells. *Slc23a2* deletion further reduced 5hmC in all cell types (**Fig. 3A-B**). Therefore, ascorbate depletion cell-autonomously inhibits TET activity in *Tet2-*deficient leukemia cells, suggesting it inhibits additional TET enzymes.

**Figure 3.**
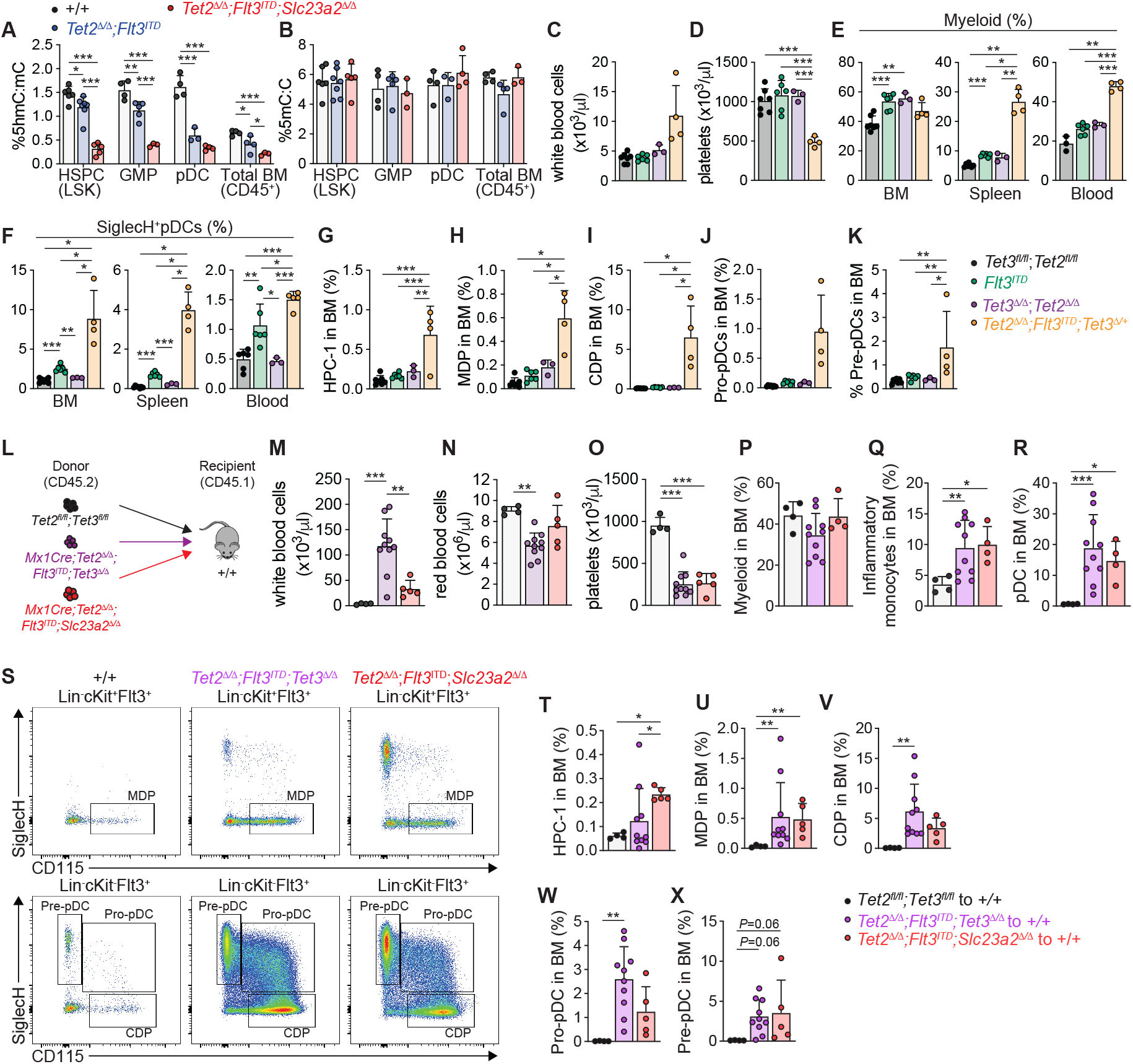
Ascorbate suppresses leukemia development by activating Tet2 and Tet3. (**A-B**) 5hmC:mC and 5mC:C ratio in sorted hematopoietic cells from the indicated genotypes. (**C-K**) Hematopoietic analysis of *Tet2^Δ/Δ^;Tet3^Δ/+^;Flt3^ITD^*mice and littermate controls 4-6 weeks after deletion. (**L-X**) Analysis of transplant recipients of *Tet2^Δ/Δ^*;*Flt3^ITD^*;*Tet3^Δ/Δ^*or *Tet2^Δ/Δ^*;*Flt3^ITD^*;*Slc23a2^Δ/Δ^*or wild-type control bone marrow. Shown are (**M-R**) mature cells; (**S**) flow plots of pDC lineage progenitors; (**T-X**) frequencies of oligopotent and pDC lineage progenitors in the bone marrow. Statistical significance was assessed with a one-way ANOVA (A-B, E, BM; G, K, M-P and V-X), Brown-Forsythe ANOVA (C-D, E, spleen and blood; F, H-J and Q-R), or a Kruskal-Wallis test (T-U).

Hematopoietic *Tet2;Tet3* deletion causes a lethal AML^49,50^. We thus tested if TET2/TET3 loss phenocopies ascorbate depletion. *Mx1Cre;Tet2^Δ/Δ^;Tet3^Δ/+^;Flt3^ITD^*mice exhibited leukocytosis, thrombocytopenia, splenomegaly, myelomonocytic and pDC expansion, and B cell depletion as compared to double or single knockout or wild-type controls (**Fig. 3C-F, Fig. S4A-E**). *Tet2^Δ/Δ^;Tet3^Δ/+^;Flt3^ITD^*mice depleted HSCs and MPPs, accumulated some oligopotent or restricted progenitors, and expanded pDC lineage progenitors (**Fig. 3G-K, Fig. S4F-H**). *Tet2^Δ/Δ^;Tet3^Δ/+^;Flt3^ITD^*phenotypes were similar to the phenotypes of *Slc23a2^Δ/Δ^;Tet2^Δ/Δ^;Flt3^ITD^*mice or of *Gulo^−/−^* mice transplanted with *Tet2^Δ/Δ^;Flt3^ITD^*cells. We transplanted *Tet2^Δ/Δ^;Tet3^Δ/Δ^;Flt3^ITD^*and *Slc23a2^Δ/Δ^;Tet2^Δ/Δ^;Flt3^ITD^* cells to recipient mice to directly compare their phenotypes (**Fig. 3L**). *Tet2^Δ/Δ^;Tet3^Δ/Δ^;Flt3^ITD^* and *Slc23a2^Δ/Δ^;Tet2^Δ/Δ^;Flt3^ITD^* donor cells had leukocytosis and thrombocytopenia, and showed similar changes in HSCs and progenitors, including a pronounced expansion of pDC lineage progenitors (**Fig. 3M-X, Fig. S4I-O**). Therefore, ascorbate depletion promotes pDC-AML by inhibiting TET2/3 activity.

### Leukemia cells take up ascorbate as a tumor suppressor response that depletes systemic ascorbate

Transformed cells induce tumor suppressor responses, such as increased p53, p16^Ink4a^, or p19^Arf^ expression, as a mechanism to limit cancer progression^51,52^. If ascorbate is a bona fide tumor suppressor, then leukemic cells could elevate it to limit leukemogenesis. Consistent with this, bulk leukemia cells accumulated intracellular ascorbate compared to normal hematopoietic cells (**Fig. 1C**). To understand if LSCs specifically accumulate ascorbate, we measured ascorbate levels in *Tet2^Δ/Δ^;Flt3^ITD^* Lin^−^Sca1^+^Kit^+^ leukemia cells isolated from wild-type transplant recipient mice. LSCs and other leukemic progenitors accumulated high levels of ascorbate compared to normal HSPCs. This was blocked by *Slc23a2* deletion (**Fig. 4A**).

**Figure 4.**
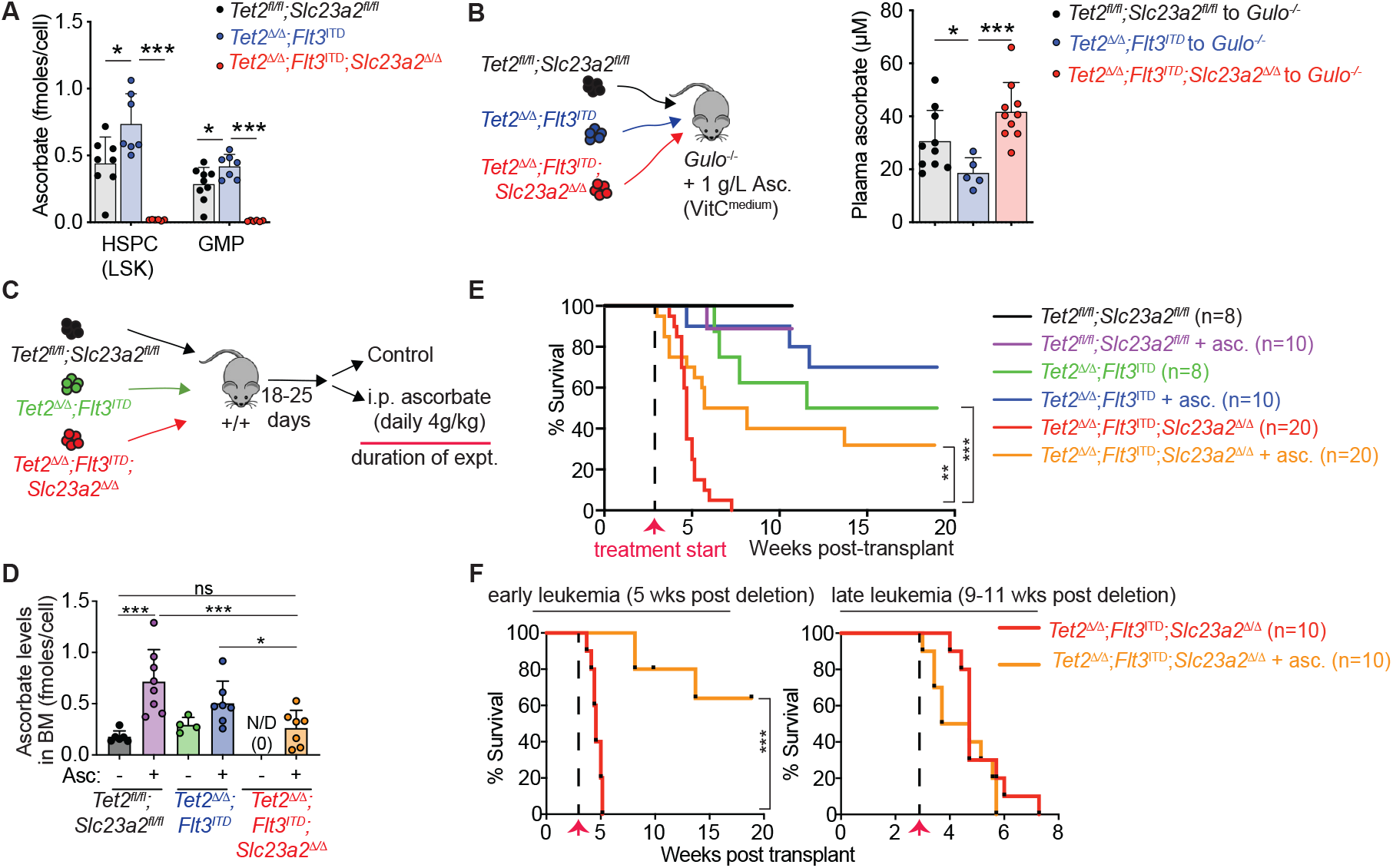
Leukemia cells accumulate ascorbate as a tumor suppressor response that depletes systemic ascorbate and restoring it blocks leukemia growth but not normal hematopoiesis. (**A-B**) Intracellular and plasma ascorbate levels from the indicated conditions. (**C**) Irradiated mice were transplanted with *Tet2^fl/fl^;Slc23a2^fl/fl^*, *Mx1Cre*;*Tet2^Δ/Δ^*;*Flt3^ITD^* or *Mx1Cre*;*Tet2^Δ/Δ^*;*Flt3^ITD^*;*Slc23a2^Δ/Δ^*bone marrow 5 weeks (early leukemia) or 9-11 weeks (late leukemia) post deletion and then treated with ascorbate. (**D**) Intracellular ascorbate levels in bone marrow cells isolated from recipient mice. (**E-F**) Kaplan-Meier survival curve of all mice (**E**) or of mice separated by transplant time (**F**). Statistical significance was assessed with Brown-Forsythe ANOVA (A-B), Kruskal-Wallis test (D), or Mantel-Cox test (E-F).

Circulating ascorbate is often depleted in leukemia patients for unknown reasons^17,30-36^. Leukemias in wild-type mice do not deplete circulating ascorbate (**Fig. 1C**), but these mice are capable of endogenous synthesis that maintains ascorbate levels, unlike GULO-deficient humans. To understand if leukemia depletes systemic ascorbate in a GULO-deficient organism, we transplanted wild-type, *Tet2^Δ/Δ^;Flt3^ITD^,* or *Slc23a2^Δ/Δ^;Tet2^Δ/Δ^;Flt3^ITD^* cells into *Gulo^−/−^*mice provided with a sufficiently high but not saturating ascorbate dietary dose (1 g/L in the water) to achieve a plasma concentration of 30 μM (**Fig. 4B**). This concentration corresponds to the ∼25^th^ percentile of concentration in the U.S. population and is well within the normal range of ∼10-100 μM^27,29^. The presence of leukemia cells depleted plasma ascorbate. Deletion of *Slc23a2* in leukemia cells to abolish their ascorbate uptake restored plasma ascorbate to wild-type levels (**Fig. 4B**). Therefore, leukemia cells siphon ascorbate from the plasma as a tumor suppressor response that inhibits leukemia progression but depletes the host of systemic ascorbate.

### Restoration of ascorbate inhibits growth of early-stage established leukemia without inhibiting normal hematopoiesis

We then tested if restoring this tumor suppressor response by repleting intracellular ascorbate can inhibit established *Slc23a2^Δ/Δ^*leukemia growth. Because *Slc23a2* is required for ascorbate uptake at physiological concentrations, we treated mice bearing established *Slc23a2^Δ/Δ^;Tet2^Δ/Δ^;Flt3^ITD^* or *Tet2^Δ/Δ^;Flt3^ITD^* leukemias with high-dose intraperitoneal ascorbate, which similar to intravenous administration transiently raises ascorbate to supraphysiological levels and produces oxidized ascorbate that can be taken up through GLUT transporters^13^, bypassing SLC23A2. Ascorbate was administered daily starting at 18-25 days after transplant, just before the onset of mortality in mice with *Slc23a2^Δ/Δ^;Tet2^Δ/Δ^;Flt3^ITD^*leukemia (**Fig. 4C**).

Parenteral ascorbate treatment restored intracellular ascorbate in *Slc23a2^Δ/Δ^* leukemia cells from undetectable (**Fig. 1C**) to the levels of untreated *Slc23a2^+/+^*leukemia cells (**Fig. 4D**). Treated *Slc23a2^Δ/Δ^* leukemia cells had lower ascorbate levels than treated *Slc23a2^+/+^* leukemia cells (**Fig. 4D**), suggesting supraphysiological ascorbate enters leukemia cells partly through SLC23A2 and partly through other transporters. Consistent with prior experiments (**Fig. 1D**), *Slc23a2* deletion in leukemia cells reduced survival of untreated mice (**Fig. 4E**). Ascorbate treatment extended survival of mice with *Slc23a2^Δ/Δ^*leukemia (**Fig. 4E**). One third of treated mice with *Slc23a2^Δ/Δ^* leukemias survived for at least 19 weeks post-transplant, the maximum duration of the experiment, in contrast to the 7-week maximum survival of untreated mice (**Fig. 4E**). The effect of ascorbate was bimodal. Ascorbate conferred a dramatic survival advantage on mice transplanted with leukemia cells 5 weeks after poly-I:C mediated gene deletion but not 9-11 weeks after deletion (**Fig. 4F**). This suggested that ascorbate was effective in early-stage, but not late-stage leukemia. Consistent with this, ascorbate administration inhibited leukocytosis, anemia, and thrombocytopenia in early-stage but not late-stage *Slc23a2^Δ/Δ^*leukemia for at least 14 weeks post-treatment (**Fig. S5A-C**). Ascorbate rescued expansion of *Slc23a2-*deficient pDCs in both early and late-stage disease suggesting that pDC reduction is not associated with a survival benefit (**Fig. S5D**). Parenteral ascorbate did not affect the frequency of HSCs and progenitor cells in wild type control mice, suggesting that ascorbate does not impair normal hematopoiesis and did not reverse the effects of *Tet2^Δ/Δ^;Flt3^ITD^* on hematopoiesis (**Fig. S5E-M**). Therefore, supraphysiological ascorbate, when provided early in disease, mitigates ascorbate-deficient leukemia without affecting ascorbate-sufficient normal or malignant hematopoiesis.

### Ascorbate deficiency promotes the growth of some but not all human AMLs

Ascorbate is severely depleted in many patients with blood neoplasms^17,30-36^. But it is unclear if ascorbate is a physiological regulator of human leukemias or other cancers *in vivo*. To test if ascorbate cell-autonomously regulates human leukemia growth, we deleted *SLC23A2* in AML cells and xenografted them to NSG mice (**Fig. 5A**). *SLC23A2* deletion significantly reduced ascorbate levels in xenografted cells (**Fig. 5B, 5F, 5M**), confirming that it is their physiological ascorbate transporter. *SLC23A2* deletion increased MV-4-11 and THP-1 cell growth in some tissues but not others, and decreased survival of THP-1 xenografted mice, suggesting accelerated leukemia progression (**Fig. 5C-L**). *SLC23A2* deletion did not affect the growth of a high-risk patient-derived myelodysplasia-related AML which had progressed after treatment (**Fig. 5M-P**), consistent with the lack of effect of ascorbate restoration in late-stage mouse leukemias (**Fig. 4F**). Therefore, physiological ascorbate levels *in vivo* suppress the growth of some but not other human AML cells.

**Figure 5.**
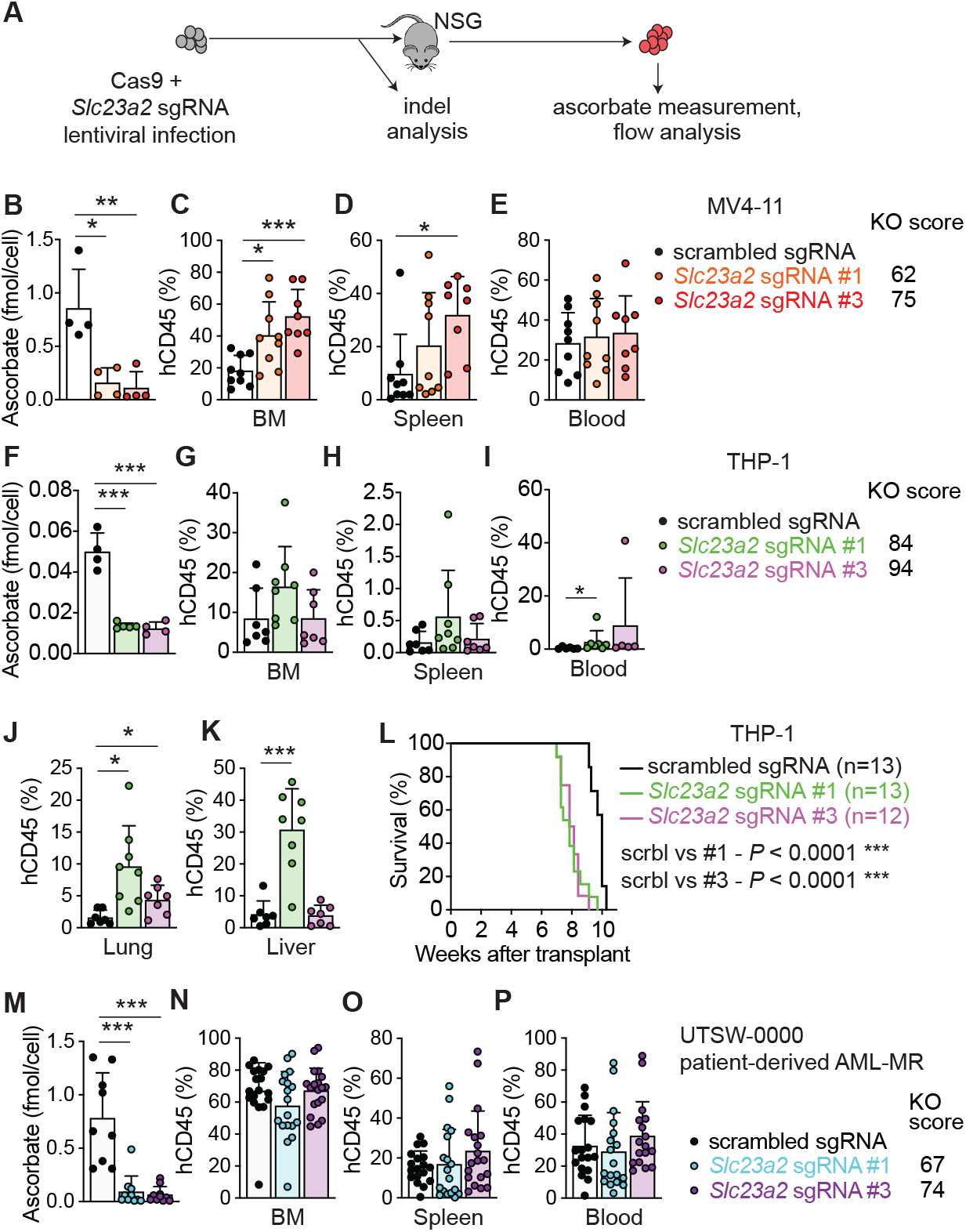
Ascorbate deficiency promotes growth of some but not all human leukemias. (**A**) Overview of experimental strategy. (**B-E**) Transplant of scrambled control or *SLC23A2*-knockout MV-4-11 cells into NSG mice. (**B**) Intracellular ascorbate levels in MV-4-11 cells sorted from NSG mice. (**C-E**) Frequency of MV-4-11 cells in NSG mice. (**F-L**) Transplant of scrambled control or *SLC23A2*-knockout THP-1 cells into NSG mice. (**F**) Intracellular ascorbate levels in THP-1 cells sorted from NSG mice. (**G-K**) Frequency of THP-1 cells in NSG mice. (**L**) Kaplan-Meier survival curve of NSG mice transplanted with THP-1 cells. (**M-P**) Transplant of CRISPR/Cas9-edited scrambled or *SLC23A2* KO cells from a patient-derived AML (myelodysplasia related, AML-MR) carrying *JAK2/TET2/WT1* mutations into NSG mice. (**M**) Intracellular ascorbate levels in AML cells sorted from NSG mice. (**N-P**) Frequency of patient-derived AML cells in NSG mice. Statistical significance was assessed with a Kruskal-Wallis test (B, D, I and M), one-way ANOVA (C, F and K), a Brown-Forsythe ANOVA (J), or a Mantel-Cox test (L).

### Ascorbate suppresses development of multiple types of leukemia

*Mx1Cre;Slc23a2^Δ/Δ^;Tet2^Δ/Δ^;Flt3^ITD^* mice showed expanded CD19^+^B220^−^ cells in the spleen at 4-6 weeks post deletion, and in the blood at 11-13 weeks (**Fig. 6A-C**). To functionally test if this indicated development of a B cell leukemia, we transplanted total bone marrow cells from *Slc23a2^Δ/Δ^;Tet2^Δ/Δ^;Flt3^ITD^*mice with either CD19^+^B220^−^ B cell neoplasm or pDC+myeloid neoplasm, or from *Tet2^Δ/Δ^;Flt3^ITD^* mice with myeloid neoplasm into sublethally irradiated recipients (**Fig. 6D**). Recipients developed blood neoplasms matching those of the donor bone marrow (**Fig. 6E-F**). Recipients of either pDC-AML or B cell *Slc23a2^Δ/Δ^;Tet2^Δ/Δ^;Flt3^ITD^*neoplastic cells developed anemia and thrombocytopenia and died earlier than recipients of *Tet2^Δ/Δ^;Flt3^ITD^* cells, consistent with our prior data that ascorbate deficiency cell-autonomously accelerates leukemogenesis (**Fig. 6G, Fig. S6A-B**). To test the leukemogenic capacity of CD19^+^B220^−^ cells, 40,000 sorted CD19^+^B220^−^ cells or 500,000 unfractionated bone marrow cells from primary recipients with B cell leukemia were transplanted into sublethally irradiated secondary recipients. Both sets of secondary recipients developed CD19^+^B220^−^ leukemia (**Fig. 6H-K, Fig. S6C**). Mice receiving purified CD19^+^B220^−^ cells had higher leukemic cell frequency than those receiving 12.5-fold more unfractionated bone marrow cells, suggesting the CD19^+^B220^−^ population contained leukemia-initiating cells (**Fig. 6I**). These data suggest that physiological ascorbate suppresses development of B cell leukemia in addition to pDC-AML.

**Figure 6.**
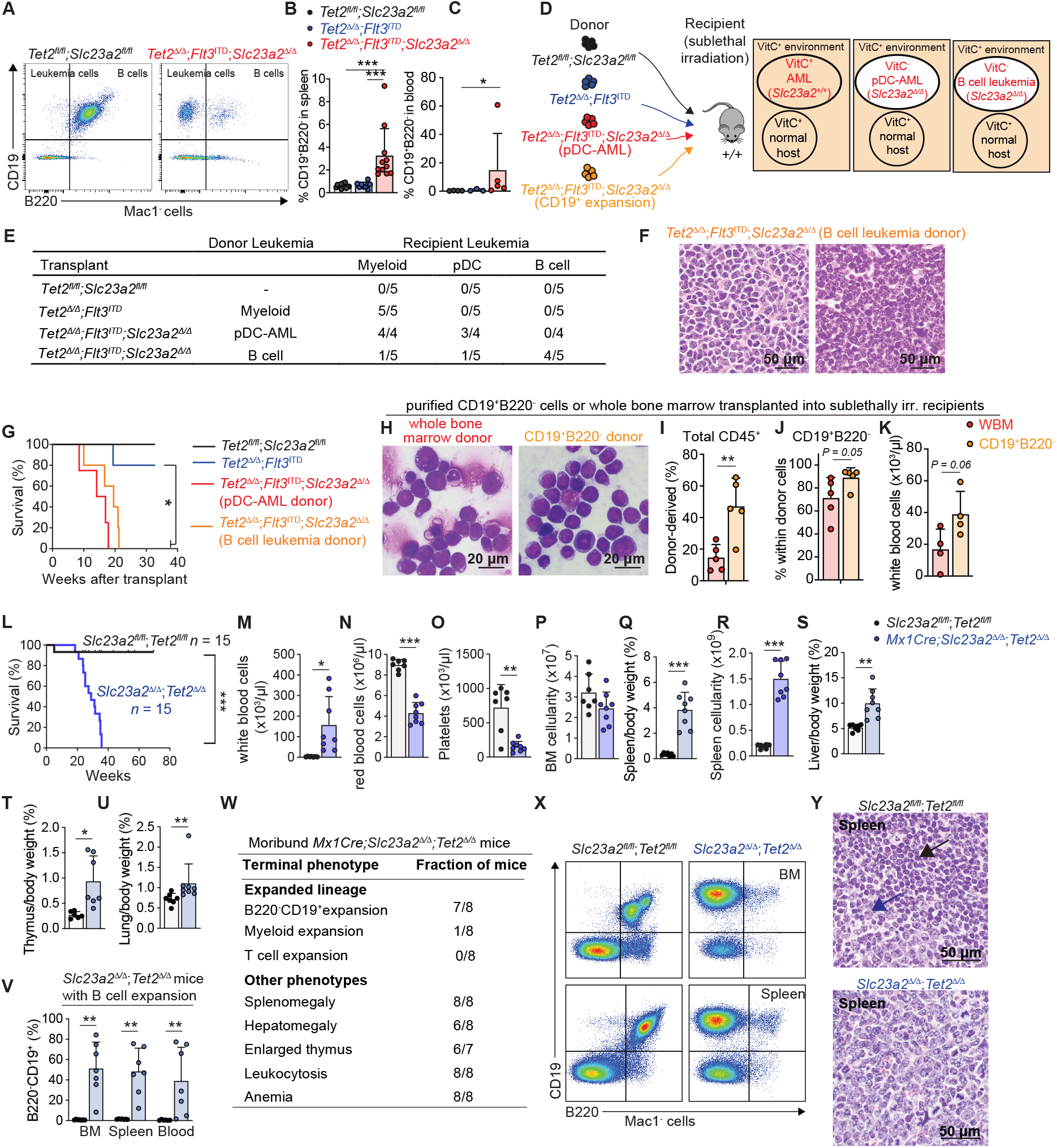
Ascorbate suppresses the development of multiple types of leukemia. (**A**) Representative flow plots of leukemic CD19^+^B220^−^ cells in the blood. (**B-C**) CD19^+^B220^−^ cell frequency in the spleen 4-6 weeks after deletion or the blood 11-13 weeks after deletion. (**D-I**) Transplantation of bone marrow from mice with pDC-AML or B cell leukemia 10 weeks after deletion into sublethally irradiated wild-type mice. (**E**) Leukemia type in recipient mice. (**F**) Representative H&E images of B-lymphoid neoplasm in liver and bone marrow of transplant recipient mice. The liver (left) and bone marrow (right) are infiltrated by sheets of medium-sized to large cells with irregular nuclei, inconspicuous to small nucleolus, and scant to moderate cytoplasm. (**G**) Kaplan-Meier survival curve of recipient mice transplanted with the indicated donor cells (n = 5 mice/genotype). (**H-K**) Transplantation of 5×10^5^ whole bone marrow cells or 4×10^4^ sorted CD19^+^B220^−^ cells from primary recipients of *Mx1Cre;Tet2^Δ/Δ^;Flt3^ITD^;Slc23a2^Δ/Δ^*cells into sublethally irradiated secondary recipients. (**H**) Representative images of bone marrow smears from mice with a B-lymphoid neoplasm. (**I-K**) Blood analysis of secondary recipients 4 weeks (**I-J**) and 8 weeks (**K**) after transplant. (**L**) Kaplan-Meier survival curve of *Mx1Cre*;*Slc23a2^Δ/Δ^*;*Tet2^Δ/Δ^*mice in weeks after deletion (n = 15 mice/genotype). (**M-V**) Hematopoietic analysis of moribund *Mx1Cre*;*Slc23a2^Δ/Δ^*;*Tet2^Δ/Δ^* mice and littermate controls. (**W**) Summary of phenotypes of moribund *Mx1Cre*;*Slc23a2^Δ/Δ^*;*Tet2^Δ/Δ^* mice. (**X-Y**) Representative flow plots and spleen H&E images in moribund *Mx1Cre*;*Slc23a2^Δ/Δ^*;*Tet2^Δ/Δ^* and control mice. Control spleen shows B-cell follicles (blue arrow, medium-sized lymphoid cells) and small lymphocytes (black arrow). In leukemic spleen, the white pulp is replaced by medium-sized blastoid lymphoid cells with vesicular chromatin, small nucleolus, and scant cytoplasm. Statistical significance was assessed with a Kruskal-Wallis test (B-C), Mantel-Cox test (G and L), t-test (I, K, N, P), Mann Whitney test (J, U and V, blood), or a Welch’s t-test (M, O, Q-T, and V, BM and spleen).

### Hematopoietic ascorbate deficiency collaborates with Tet2 loss to cause leukemia

Somatic *TET2* loss of function mutations cause clonal hematopoiesis and increase risk of leukemia, raising the question of whether ascorbate depletion modulates *Tet2*-deficient leukemogenesis. Consistent with this idea, ascorbate depletion increases *Tet2^+/−^* myeloid neoplasia^19^. To test if hematopoietic ascorbate depletion can transform *Tet2^Δ/Δ^* cells, we analyzed *Mx1Cre;Slc23a2^Δ/Δ^;Tet2^Δ/Δ^*mice. *Mx1Cre*;*Tet2^Δ/Δ^* mice develop predominantly myeloid leukemia, and less frequently B cell leukemia, with a long latency and median survival of about 80 weeks^50,53^. *Mx1Cre;Slc23a2^Δ/Δ^;Tet2^Δ/Δ^*mice developed highly accelerated leukemia with a median survival of 29 weeks after deletion (**Fig. 6L**). Moribund mice had leukocytosis, anemia, thrombocytopenia, and enlarged liver, spleen, thymus, and lungs (**Fig. 6M-W**). Analysis of moribund mice revealed CD19^+^B220^−^ B cell leukemia infiltrating the bone marrow, spleen, and blood in 7/8 analyzed mice and myeloid leukemia in 1/8 (**Fig. 6V-Y**). Therefore, hematopoietic ascorbate deficiency collaborates with TET2 deficiency to produce predominantly B cell leukemia.

### Hematopoietic ascorbate deficiency is sufficient to cause leukemia

To test whether ascorbate deficiency alone is sufficient to initiate leukemia in the absence of other engineered mutations, we aged *Mx1Cre;Slc23a2^Δ/Δ^* mice and littermate controls. *Slc23a2^Δ/Δ^* mice spontaneously developed leukemia. Moribund *Slc23a2^Δ/Δ^* mice had splenomegaly, bone marrow attrition, enlarged thymus and lungs, leukocytosis, anemia, and thrombocytopenia, consistent with development of leukemia histologically confirmed by a hematopathologist (**Fig. 7A-J**). Moribund *Slc23a2^Δ/Δ^* mice exhibited a range of blood cancers, with 5/9 having B220^−^ CD19^+^ B cell leukemia and 4/9 myeloid leukemia. In all cases, leukemia cells were present in the spleen, bone marrow and blood. *Slc23a2^Δ/Δ^* mice succumbed to disease with a median survival of 46 weeks post-deletion (**Fig. 7K-P**).

**Figure 7.**
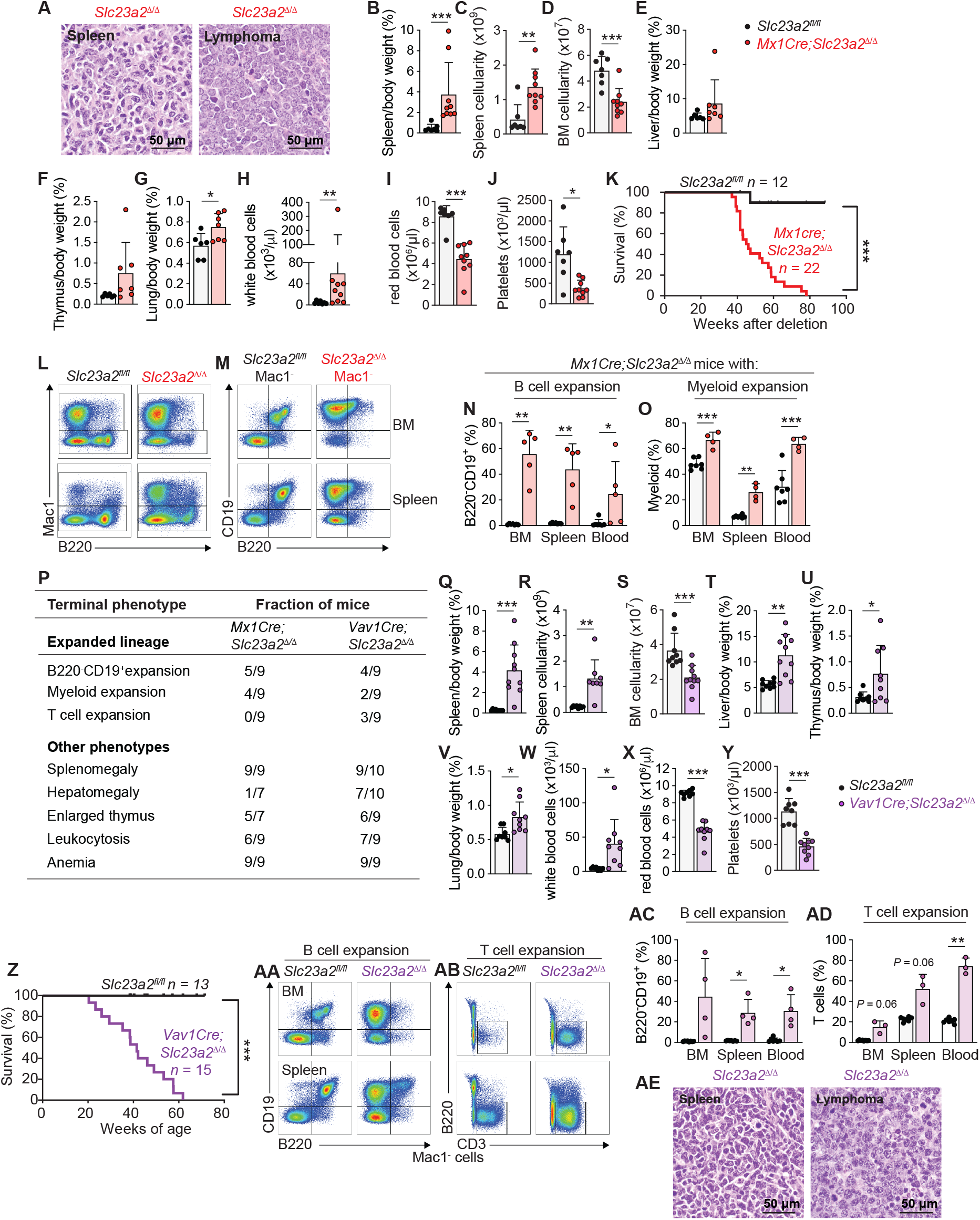
Hematopoietic ascorbate deficiency is sufficient to induce leukemia. (**A**) Representative H&E images of myeloid neoplasm in spleen and lymphoma from *Mx1Cre*;*Slc23a2^Δ/Δ^* mice. The white pulp in the spleen is completed replaced by medium-sized to large cells with irregular nuclei, variable nucleolus, and moderate cytoplasm (left). The lymphoma is filled with medium-sized to large blastoid cells with vesicular chromatin, inconspicuous to small nucleolus, and scant cytoplasm (right). (**B-J**) Relative organ weights, BM and spleen cellularity and blood analysis in moribund *Mx1Cre*;*Slc23a2^Δ/Δ^* and littermate control mice. (**K**) Kaplan-Meier survival curve of *Mx1Cre*;*Slc23a2^Δ/Δ^*mice. (**L-O**) Representative flow plots and frequencies of expanded cells in moribund *Mx1Cre*;*Slc23a2^Δ/Δ^* mice with B cell or myeloid cell expansion. (**P**) Phenotypes of moribund *Mx1Cre*;*Slc23a2^Δ/Δ^* and *Vav1Cre*;*Slc23a2^Δ/Δ^* mice. (**Q-Y**) Relative organ weights, cellularity of BM and spleen, and blood analysis in moribund *Vav1Cre*;*Slc23a2^Δ/Δ^* and littermate control mice. (**Z**) Kaplan-Meier survival curve of *Slc23a2^fl/fl^* (n = 13 mice) and *Vav1Cre*;*Slc23a2^Δ/Δ^* mice (n = 15 mice). (**AA-AD**) Flow plots and frequencies of B and T cell leukemia in bone marrow and spleen of moribund *Vav1Cre*;*Slc23a2^Δ/Δ^* and control mice with B or T cell expansions respectively. (**AE**) Representative H&E images of T cell neoplasm in spleen and lymphoma from *Vav1Cre*;*Slc23a2^Δ/Δ^* mice showing medium-sized to large blastoid lymphoid cells. Statistical significance was assessed with a Mann-Whitney test (B-C, E, H-I, and N, blood), t-test (D, G, O, BM and blood; S and Y), Welch’s t-test (F, J, N, BM and spleen; O, spleen; Q-R, T-X and AC-AD), or a Mantel-Cox test (K and Z).

To avoid any confounding effects of poly I:C-induced inflammation or deletion in other tissues in the *Mx1Cre* model, we used *Vav1Cre* to delete *Slc23a2* from fetal hematopoiesis. *Vav1Cre;Slc23a2^Δ/Δ^*mice developed leukemia with a median survival of 42 weeks of age. Moribund *Vav1Cre;Slc23a2^Δ/Δ^* mice exhibited a range of blood cancers, including myeloid, B, and T cell neoplasms (**Fig. 7P-AE**). Therefore, hematopoietic-specific ascorbate deficiency is sufficient to cause blood cancer.

## DISCUSSION

Our results nominate ascorbate as the first, to our knowledge, tumor suppressor metabolite. A tumor suppressor metabolite should meet two criteria: its depletion within cancer-initiating cells should collaborate with cancer-driver mutations to promote cancer formation; and it should be depleted in some patients with the corresponding cancer. Ascorbate fulfills these criteria: ascorbate depletion within hematopoietic cells collaborates with *Tet2^Δ/Δ^;Flt3^ITD^*to accelerate leukemogenesis, with *Tet2^Δ/Δ^* to cause leukemia, and it is sufficient to cause leukemia with high penetrance in the absence of other engineered mutations; it suppresses AML by direct activation of TET enzymes within leukemia-initiating HSPCs; and this natural tumor suppressor mechanism is disabled in leukemia patients with systemic ascorbate depletion^17,30-36^. Mutations in *SLC23A2* in leukemias have not been reported. This may be because ascorbate is depleted systemically or because some human leukemias express SLC23A1^54^, which can compensate for SLC23A2 loss. SLC23A2 is however repressed or mis-spliced in some myeloid and lymphoid leukemia cells^34,55,56^.

Ascorbate has been proposed to restrain cancer development by acting in immune, stromal, or cancer cells. However, this was not directly tested by cell-type specific manipulations *in vivo*. Conditional *Slc23a2* deletion and transplantation of purified cells suggested that ascorbate suppresses leukemogenesis by acting within leukemia-initiating cells. Pathologies such as thrombocytopenia, anemia, and shortened mouse survival develop after transplantation of *Slc23a2^Δ/Δ^* leukemia cells to mice in the presence of wild-type hematopoietic cells, indicating they are caused by *Slc23a2^Δ/Δ^*leukemia rather than by impaired *Slc23a2^Δ/Δ^* hematopoiesis. Ascorbate is a direct TET activator^48^ and its systemic depletion inhibits primarily TET2 in HSCs and multiple TET enzymes in myeloid progenitors or AML^24^. Our results with compound *Slc23a2, Tet2,* and *Tet3* knockout mice suggest TET2/TET3 are the key mediators of the action of ascorbate in AML suppression. This is consistent with findings that ascorbate suppresses *Tet2^Δ/Δ^* HSC self-renewal *in vitro* via TET3^15^ and promotes plasma cell development via TET2/TET3^57,58^. In addition to AML patients, ascorbate depletion is also prevalent in patients with B cell neoplasms, and supraphysiological ascorbate suppresses B cell lymphoma growth by promoting TET activity^14^. *TET2* mutations are present in B cell neoplasms^59^ and *Tet2* deletion promotes B cell leukemia in mice^60^, including of a CD19^+^B220^−^ neoplasm^61^. *Tet2;Tet3* deletion causes myeloid^49^, B cell^62^, and T cell^63^ malignancies. Therefore, TET2/TET3 inhibition may contribute to development of multiple neoplasms in *Slc23a2^Δ/Δ^*mice.

Our findings reveal pDC-AML developmental origins in mice. TET deficiency collaborates with FLT3 activation to expand early pDC-lineage progenitors. We propose that pDC expansion may be an unrecognized feature of other AML models with B220^+^ cell expansion^64,65^. pDC-AML has a poor prognosis compared to other AML types. Our results that the LSC in pDC-AML is contained within multipotent stem and progenitor cells and not pDC lineage cells are consistent with prior results that CD34^+^ blasts but not pDCs can transplant human pDC-AML to immunocompromised mice^46^. Our findings nominate ascorbate as a candidate therapeutic to inhibit aberrant self-renewal and promote differentiation of leukemic HSPCs in myeloid neoplasms with pDC expansions.

Dietary molecules have complex effects in the body. If their cellular and molecular locus of action within a complex tissue is unclear, then prospects for generalizing and applying any discoveries will be limited. Ascorbate is a prime example of this. Oral ascorbate supplementation can raise plasma concentrations up to the high end of the physiological range (∼80-100 μM). However, most ascorbate supplementation trials in cancer patients did not consider the cancers, cell types, and molecules that physiological ascorbate acts on^66-70^. Treatment trials using oral supplementation focused on advanced solid cancers and specifically excluded leukemia patients^66,67^, even though there was little evidence for a physiological role of ascorbate in advanced solid cancers. Prevention trials^68-71^ did not select for individuals with baseline ascorbate deficiency, even though oral supplementation has at most modest effects on ascorbate concentration in individuals who are baseline-sufficient^72,73^.

Ascorbate can promote TET activity^74^ and improve outcomes in some AML patients^75^. The recent EVITA phase 2 randomized trial found that ascorbate supplementation of patients with clonal cytopenia or early-stage myeloid neoplasms who had below-average concentrations at baseline increased survival^37^. Our findings provide mechanistic support for ascorbate repletion trials in patients with early-stage myeloid or B cell neoplasms who are ascorbate-deficient at baseline.

The discovery that ascorbate is a natural tumor-suppressor metabolite raises the question of whether there are other such metabolites. Supraphysiological elevation of some metabolites inhibits cancer development. In addition to ascorbate^5,8^, these include β-hydroxybutyrate^2^, ursodeoxycholate^76,77^, and α-ketoglutarate^3,4,78^. Conversely systemic depletion of some metabolites promotes cancer development. In addition to ascorbate^8,24^, these include choline-methionine depletion in hepatocellular carcinoma^79^, and folate depletion in some colorectal cancers^80^. Such metabolites may also serve as tumor suppressors if they are depleted in patients, and if their depletion within cancer-initiating cells promotes or is sufficient to cause development of cancer.

## Methods

### Mice

*Tet2^fl/fl^* (JAX:017573)^81^, *Tet3^fl/fl^* (JAX:031015)^82^, *Flt3^ITD^* (JAX:011112)^83^, *Slc23a2^−/−^* (JAX:027439)^39^, *Slc23a2^fl/fl^*^41^, *Gulo^−/−^* (JAX:004317)^84^, *Mx1-cre* (JAX:003556)^85^, *Vav1Cre^86^* and NSG (JAX:005557)^87^ mice were previously described. Except for NSGs, mice were on a C57BL background. Both male and female mice were used in all studies. Mice were injected every other day with 40 μg poly (I:C) (Cytiva) for a total of five intraperitoneal injections starting at 4-6 weeks of age to induce Cre expression. Mice were analyzed 4-6 weeks after the first day of poly (I:C) injection, or as indicated in the text. C57BL/Ka-Thy-1.1 (CD45.2) and C57BL/Ka-Thy-1.2 (CD45.1) mice were used for transplantation experiments. *Gulo*^−/−^;CD45.1 recipient mice were generated by crossing *Gulo*^−/−^ mice with CD45.1 mice. *Gulo^−/−^*mice were provided with 1% ascorbate diet (Envigo TD.160622) for breeding, maintenance, or repletion experiments, or 1 g/L ascorbate in the drinking water changed every two days to provide sufficient ascorbate, or 50-100 mg/L ascorbate in the drinking water to test the effects of chronic ascorbate deficiency, or no supplementary ascorbate for short-term complete deprivation experiments as described in the text. For high dose ascorbate treatment experiments, mice received daily IP injections of ascorbate (4 g/kg), 5 days per week, starting at 18-25 days after transplant. Mice were housed in the Animal Resource Center of UT Southwestern and procedures were approved by the UT Southwestern Institutional Animal Care and Use committee.

### Hematopoietic or leukemia cell isolation and analysis

Bone marrow cells were obtained by flushing femurs with a 25G needle, or crushing femurs, tibias, vertebrae, and pelvic bones with a mortar and pestle in staining medium consisting of Ca^2+^/Mg^2+^-free Hank’s balanced salt solution (HBSS; Gibco), supplemented with 2% heat-inactivated bovine serum (Gibco). Spleens were mechanically dissociated by trituration in staining medium. Livers were enzymatically digested for 30 minutes at 37°C in 1.5 ml RPMI-1640 (Sigma), containing 250 μg/ml liberase (Roche) and 100 μg/ml DNase I (Roche). Cell suspensions were filtered through a 40 μm strainer. Cell number was measured with a Vi-CELL cell viability analyzer (Beckman Coulter). Blood was collected by cardiac puncture using a 25G needle and mixed in a tube containing 5 μl 0.5M EDTA. Complete blood cell counts were determined using a Hemavet HV950 (Drew Scientific) or Element HT5 (Heska). For hematopoietic analysis, 40 μl blood was lysed in 1 ml of ammonium chloride buffer (155mM NH_4_Cl; 10 mM KHCO_3_; 0.1 mM EDTA) for 10-15 minutes at 4°C. Cells were incubated with fluorescently conjugated antibodies for 90 minutes on ice when using CD34 or for 30 minutes at 4°C. Cells were washed with staining medium and resuspended in staining medium containing 1 μg/ml DAPI or 1 μg/ml propidium iodide for live/dead discrimination. Cell populations were defined as previously described^47,88-95^ with the following markers: CD150^+^CD48^−^Lineage^−^Sca-1^+^Kit^+^ hematopoietic stem cells (HSCs), CD150^−^CD48^−^Lineage^−^Sca-1^+^Kit^+^ multipotent progenitor cells (MPPs), CD150^−^CD48^+^Lineage^−^Sca-1^+^Kit^+^ hematopoietic progenitor cells (HPC-1), CD150^+^CD48^+^Lineage-Sca-1^+^Kit^+^ hematopoietic progenitor cells (HPC-2), CD34^+^CD16/32^−^ Lineage^−^Sca-1^−^Kit^+^ common myeloid progenitors (CMPs), CD34^+^CD16/32^+^Lineage^−^Sca-1^−^Kit^+^ granulocyte–monocyte progenitors (GMPs), CD34^−^CD16/32^−^Lineage^−^Sca-1^−^Kit^+^ megakaryocyte– erythroid progenitors (MEPs), Lineage^−^Sca-1^+^Kit^+^ multipotent/oligopotent progenitors, Mac-1^+^ myeloid cells, Mac1^+^CD115^+^Ly6C^+^Ly6G^−^ inflammatory monocytes, Mac1^+^CD115^−^ Ly6C^mid^Ly6G^+^ neutrophils, Mac1^−^CD3^+^ T cells, Mac1^−^B220^+^CD19^+^ B cells, Mac1^−^B220^+^CD19^−^ leukemic cells, Mac1^−^CD19^+^B220^−^ leukemic cells, Mac1^−^B220^+^CD19^−^SiglecH^−^PDCA^+^, Mac1^−^ B220^+^CD19^−^SiglecH^+^PDCA^+^, or Mac1^−^B220^−^CD19^−^SiglecH^+^PDCA^+^ plasmacytoid dendritic cells (pDCs), Lineage^−^Kit^+^Sca-1^−^Flt3^+^CD115^+^SiglecH^−^ macrophage-dendritic cell progenitors (MDPs), Lineage^−^Kit^−^CD115^+^Flt3^+^SiglecH^−^ common dendritic cell progenitors (CDPs), Lineage^−^Kit^−^ Flt3^+^CD115^+^SiglecH^+^ pro-plasmacytoid dendritic cells (pro-pDCs), and Lineage^−^Kit^−^Flt3^+^CD115^−^ SiglecH^+^ pre-plasmacytoid dendritic cells (pre-pDCs). The Lineage cocktail for HSCs and progenitors consisted of CD2, CD3, CD5, CD8, Ter119, B220, and Gr-1 antibodies. For flow cytometry experiments, combinations of antibodies were used against the following cell-surface markers, conjugated to Violet Fluor 450, Brilliant Violet 421, Brilliant Violet 510, FITC, PE, PE594, PerCP-Cy5.5, PE-Cy7, APC, eFluor 660, Alexa Fluor 700, RedFluor 710, APC/Fire 750, or APC-Cy7: CD2 (RM2-5, Tonbo Biosciences/BioLegend), CD3 (17A2, Tonbo Biosciences/BioLegend), CD5 (53-7.3, BioLegend), CD8a (53-6.7, Tonbo Biosciences/BioLegend), CD11b (Mac1) (M1/70, Tonbo Biosciences/BioLegend), CD16/32 (93, BioLegend), CD34 (RAM34, Invitrogen), CD45.1 (A20, Tonbo Biosciences), CD45.2 (104, Tonbo Biosciences/BioLegend), CD45R (B220) (RA.3-6B2, Tonbo Biosciences/BioLegend), CD48 (HM48-1, BioLegend), CD115 (CSF-1R) (AFS98, BioLegend), CD117 (Kit) (2B8, BioLegend), CD135 (Flt3) (A2F10, Invitrogen), CD150 (TC15-12F12.2, BioLegend), CD317 (PDCA-1) (927, BioLegend), Ly-6A/E (Sca-1) (D7, BioLegend), Ly-6C (HK1.4, BioLegend), Ly-6G (1A8, BioLegend), Gr-1 (RB6-8C5, Tonbo Biosciences or Biolegend); SiglecH (551, BioLegend). Analysis and sorting were performed using the FACSAria flow cytometer (BD Biosciences) or a FACSCanto (BD Biosciences). Data were analyzed using FlowJo (Flowjo LLC) or FACSDiva (BD Biosciences).

### Bone marrow and AML transplantation assays

Recipient mice (CD45.1) were irradiated using an XRAD 320 X-ray irradiator (Precision X-Ray) with two doses of 5.4 Gy (10.8 Gy total) delivered at least 3 hours apart (for lethal irradiation) or with 1 dose of 5.4 Gy (50% of lethal dose) or 7.56 Gy (70% of lethal dose). For non-competitive transplants, 3-5 × 10^6^ bone marrow donor (CD45.2) cells were transplanted into the retro-orbital venous sinus of anesthetized recipient (CD45.1) mice. For transplants of leukemia cells in competition with normal hematopoietic cells, 4 × 10^4^ bone marrow donor (CD45.2) cells from *Tet2^fl/fl^;Slc23a2^fl/fl^*, *Mx1Cre*;*Tet2^Δ/Δ^*;*Flt3^ITD^*or *Mx1Cre*;*Tet2^Δ/Δ^*;*Flt3^ITD^*;*Slc23a2^Δ/Δ^* mice were mixed with 3.6 × 10^5^ wild-type competitor (CD45.1) cells and transplanted into recipient (CD45.1/CD45.2) mice. Transplantation of sorted cell populations was performed as indicated in the text. Recipient mice were maintained on antibiotic water (Baytril 0.08 mg/ml) for 1 week pre-transplantation, and for 4 weeks after transplantation. Blood was obtained from the tail veins of recipient mice every four weeks up to 16 weeks after transplantation, or as indicated in the text. Red blood cells were lysed in ammonium chloride lysis buffer. The remaining cells were stained with antibodies against CD45.2, CD45.1, CD11b (Mac1), CD45R (B220), CD19, SiglecH, CD317 (PDCA-1), and CD3 and analyzed by flow cytometry. For transplantation of human AML cells, NSG mice were irradiated with one dose of 2.25 Gy and 1 × 10^6^ MV-4-11, 2 × 10^6^ THP-1, or 5 × 10^4^ patient-derived AML cells were transplanted by injection into the retro-orbital venous sinus or the tail vein of anesthetized recipients 1 to 24 hours post irradiation. Recipient mice were maintained on antibiotic water (Amoxicillin 0.25 mg/ml) during the transplant. Bone marrow cells for analysis of transplant recipient mice were obtained by crushing femurs, tibias, vertebrae, and pelvic bones.

### Ascorbate measurement

Ascorbate was extracted from hematopoietic cells or plasma immediately after isolation using ice-cold 25:25:50 methanol:acetonitrile:water (v:v:v) + 10 mM EDTA or 10% metaphosphoric acid, with ^13^C-ascorbate as an internal standard. For measurements in purified human AML cells, 2×10^5^ human CD45^+^ cells were sorted from bone marrow of recipient NSG mice into 200 μl Ca^2+^/Mg^2+^-free HBSS supplemented with 2% heat-inactivated bovine serum, washed twice with ice-cold PBS and then resuspended 1:1 with 40 μl 10% metaphosphoric acid with 1 mM EDTA and 100 pmoles ^13^C-ascorbate. Samples were vortexed for 1 minute and then centrifuged at 20,000 g for 15 minutes at 4°C. Supernatant from sorted cells and plasma was transferred to a new tube and stored at –80°C until analysis. Ascorbate levels were quantified by liquid chromatography-tandem mass spectrometry (LC-MS/MS) using an AB Sciex 6500+ QTrap mass spectrometer operating in negative mode coupled to a Shimadzu UHPLC system with a Scherzo SM-C18 column (Imtakt) to detect the transitions 175/115 (ascorbate) and 176/116 (13C-ascorbate) as previously described^24^. Data analysis was performed with MultiQuant (SCIEX).

### Measurement of 5hmC, 5mC and C by LC-MS/MS

To measure modified cytosines in purified populations, cells were sorted by flow cytometry into Ca^2+^/Mg^2+^-free HBSS supplemented with 2% heat-inactivated bovine serum, washed twice with ice-cold PBS, and the genomic DNA was purified using Quick-DNA Miniprep kit (Zymo) according to the manufacturer’s instructions with the following modifications. Lysis buffer was supplemented with 2.4U/sample of proteinase K (NEB) and 0.2 mg/ml RNase A (NEB) and samples were eluted in 2 × 11 μl H_2_O containing 40 μM butylated hydroxytoluene (BHT, Sigma). Total genomic DNA from sorted cells was digested for 3 hours at 37°C with 7.5U/sample DNA Degradase Plus (Zymo), 10U/sample micrococcal nuclease (Thermo Scientific) and the following internal standards added to the mix: ^13^C,^15^N2-2-deoxycytidine (^13^C,^15^N2-^2^dC), D3-5-methyl-2-deoxycytidine (D3-5mdC), D3-5-hydroxymethyl-2-deoxycytidine (D3-5hmdC) (Toronto Research Chemicals). Samples were then filtered with a Nanosep 3k column to remove enzymes, and filtrates were analyzed by LC–MS/MS. For 5mC and 5hmC, liquid chromatography was carried out with a Scherzo SM-C18 (Imtakt) reverse phase C18 column, 3 μm, 2.1× 100 mm, or with an F5 reverse phase column coupled to an AB Sciex QTRAP 6500+ or 7500 operating in MRM mode as previously described^26^. For the SM-C18 column, chromatographic separation was performed using a flow rate of 0.5 mL/min. The mobile phases consisted of LC-MS-grade water containing 0.1% (v/v) formic acid (mobile phase A) and LC-MS-grade acetonitrile containing 0.1% (v/v) formic acid (mobile phase B). The gradient started at 0% B from 0–2.5 min, followed by a linear increase to 100% B from 2.5–24 min. The gradient was maintained at 100% B until 25 min, then decreased to 0% B at 27 min and maintained at 0% B until the end of the run at 30 min. For the F5 reverse phase column, chromatographic separation was performed using a flow rate of 0.25 mL/min. The mobile phases consisted of LC-MS-grade water containing 0.1% (v/v) formic acid (mobile phase A) and LC-MS-grade acetonitrile containing 0.1% (v/v) formic acid (mobile phase B). The gradient elution program was as follows: 0% B was maintained from 0 to 3 min, followed by a linear increase to 95% B over 3-24 min. The column was maintained at 95% B from 24 to 26 min. At 26.2 min, the mobile phase composition was rapidly returned to 0% B and maintained until the end of the run at 30 min. Data analysis was performed with MultiQuant (SCIEX).

### Histology and immunofluorescence

Bone marrow, spleen, liver and lymphoid tissue from primary or transplant recipient mice were collected and fixed in 10% formalin for at least 24 hours. After fixation, tissues were submitted to the UTSW Histo Pathology Core or UTSW Tissue Management Shared Resource for processing, paraffin embedding, sectioning and hematoxylin & eosin staining. Bone marrow smears were collected from transplant recipient mice, fixed and stained with Giemsa (Sigma-Aldrich). Marrow smears and H&E sections of bone marrow, spleen, and liver tissue were evaluated for tissue architecture, tumor cell infiltration and hematopoiesis by a pathologist (W.C.). For immunofluorescence staining, whole femurs were dissected, cut in a sagittal section and fixed with 4% PFA at 4°C for 20 minutes in a 96-well plate. Femurs were washed with PBS for 15 minutes and blocked in 10% donkey serum in Ca^2+^/Mg^2+^-free HBSS for 1 hour at room temperature. Femurs were stained for 2 hours at 4°C in the dark with a combination of the following antibodies: SiglecH-PE (BioLegend, 129606, 1/40), CD45R(B220)-APC (Tonbo Biosciences, 20-0452, 1/40), CD317 (PDCA-1) (Abcam, ab246508, 1/80). Femurs were washed 3x with PBS for 15 minutes and then stained for 1 hour at room temperature in the dark with the following antibodies: rabbit Alexa Fluor 488 (Invitrogen, A32790; 1/150) and Hoechst (Invitrogen, H3570, 1/1000). Femurs were washed 3x with PBS for 10 minutes and then imaged cut-face down in a coverslip-bottom well containing PBS. Images were acquired on a Zeiss LSM780 inverted confocal microscope using the tiling function and tiled images were then stitched together using Zen software (Zeiss) to create a composite image of the tissue.

### Human AML cells

MV-4-11 cells (a gift from C.C. Zhang) and THP-1 cells (a gift from S. Chung) were cultured in 90% RPMI (ATCC), supplemented with 10% FBS and 100U/mL penicillin-streptomycin. Patient-derived AML cells had mutations in *JAK2/TET2/WT1* and were from a patient with MDS/MPN treated with Azacytidine and Ruxolitinib with progression to AML at the time of bone marrow sampling. The sample was obtained under an IRB to S. Chung. Based on AML progression from MDS/MPN and failure of azacytidine treatment, this is clinically classified as a high-risk AML. Patient AML cells were transplanted two times in NSG mice, freeze/thawed, and cultured in StemSpan SFEM II (Stem Cell Technologies) supplemented with 100 ng/ml human FLT3-ligand (Peprotech), 100 ng/ml human stem cell factor (SCF, Peprotech), 50 ng/ml human thrombopoietin (TPO, Peprotech), 20 ng/ml human interleukin 3 (IL-3, Peprotech), and 20 ng/ml human IL-6 (Peprotech). HEK293T cells were purchased from ATCC (CRL-3216) and maintained in 90% D-MEM supplemented with 10% heat-inactivated FBS and 100U/mL penicillin-streptomycin.

### CRISPR/Cas9-mediated generation of knockout cells

A lentivirus vector, pLentiCRISPRv2 (Addgene, 52961)^96^, containing hSpCas9 was used to generate *SLC23A2-*deficient leukemia cells. For each knockout cell line, five sgRNAs targeting the *SLC23A2* gene were designed using the Genetic Perturbation Platform (GPP) sgRNA designer (Broad Institute). Cloning of the sgRNAs into the pLentiCRISPRv2 vector was performed according to instructions on the GPP web portal. For virus production, 8 million HEK293T cells were seeded per 100-mm dish 1 day prior to transfection. pLentiCRISPRv2 vector (5 μg) containing either scrambled or a sgRNA targeting the *SLC23A2* gene was co-transfected with 5 μg packaging plasmid psPAX2 (Addgene, 12260) and 2 μg envelope plasmid pMD2.G (Addgene, 12259) in HEK293T cells using Lipofectamine 2000 (Invitrogen) according to the manufacturer’s instructions. After 4 to 6 hours, the medium was changed to DMEM supplemented with 30% heat-inactivated FBS and 100 U/mL penicillin-streptomycin. Medium containing virus particles was harvested at 48 hours after transfection and filtered through a 0.45-μm membrane. MV-4-11, THP-1, or patient-derived AML cells were transduced with lentivirus in the presence of 8 μg/mL polybrene by spinfection at 2,000 xg at 30°C for 2 hours. Medium containing virus particles was removed after 3 hours and changed to regular cell culture media. After 72 hours, infected cells were subjected to puromycin selection (1 μg/mL for MV-4-11 and patient-derived AML, and 2 μg/mL for THP-1 cells) for at least 4 days, or until control cells without virus died. For knockout efficiency analysis genomic DNA was extracted using the Quick DNA mini prep kit (Zymo) according to the manufacturer’s instructions. DNA surrounding the sgRNA targeting sites was amplified by PCR and the product was purified with DNA clean and concentrator kit (Zymo) before Sanger sequencing (UTSW McDermott Center). Knock-out scores were calculated from sequencing chromatograms using ICE analysis (Synthego). For each cell line, two sgRNAs that yielded the highest knock-out efficiency were selected to knock out *SLC23A2* for transplantation experiments. Selected sgRNA sequences: scrambled, GAGGCTAAGCGTCGCAA; *SLC23A2* sgRNA#1, AAGACAGGATGGCTCGAGCA; *SLC23A2* sgRNA#3, GTGACTACTACGCCTGTGCA. PCR primers to amplify region surrounding sgRNA target sites: *SLC23A2* sgRNA#1 forward, GTGGCAATGGGGAGGATGAA; *SLC23A2*sgRNA#1 reverse GATGCTGACGCAGCAAAGAG; *SLC23A2*sgRNA#3 forward, CTGGTCCTGGTCACTTTCGC; *SLC23A2* sgRNA#3 reverse, AGGAATTCCGTTCCACGTCAT. Sequencing primers: *SLC23A2* sgRNA#1, GAGGATGAAGAAGCCACTG; *SLC23A2* sgRNA#3, ACAAAGGCAGTGGTGTATC.

### Statistical analysis

Most figure panels show the pooled results from mice analyzed in multiple independent experiments. Prior to analyzing the statistical significance of differences among treatments, we tested if the data was normally distributed and if variance significantly differed among treatments. To test for normal distribution, we performed the Shapiro-Wilk test when 3 ≤ n < 20 or the D’Agostino & Pearson test when n ≥ 20. To test for differences in variance we performed F-tests. If the data did not significantly deviate from normality (p < 0.01 for at least one treatment), we used a parametric test and if they did, a non-parametric test. To assess statistical significance of differences in means between treatments, we used a *t*-test or one-way ANOVA for data that was normally distributed and did not have significantly different variances, or a *t*-test with Welch’s correction or Brown-Forsythe ANOVA for data that was normally distributed and had significantly different variances, or a Mann-Whitney or Kruskal-Wallis test for data that was not normally distributed. To assess statistical significance of differences in survival between treatments we used a Mantel-Cox test. Results were graphed and statistical analysis performed using GraphPad Prism v11.

## Acknowledgements

The work was funded by grants from the Cancer Prevention and Research Institute of Texas (CPRIT) (RR180007, RP250280, RP260427), American Society of Hematology (Faculty Scholar award), Moody Foundation, Alex’s Lemonade Stand Foundation (‘A’ Award), an American Cancer Society and Simmons Cancer Center Institutional Research grant (IRG-17-174-13, P30CA142543), the Haggerty Foundation, the Rally Foundation (25IN22), and the National Institutes of Health (R01DK125713, R01HL161387) to MA. We thank Tom Mathews, Lauren Zacharias, and the Children’s Research Institute (CRI) Metabolomics facility (supported by the CPRIT Core Facilities Support Award RP240494 to Ralf DeBerardinis) for mass spectrometry support, the UTSW Histo Pathology Core and UTSW Tissue Management Shared Resource for sectioning and H&E staining, the North Texas Clinical Pharmacology Cancer Core (supported by the CPRIT Core Facilities Support Award RP210209 to LL) for 5hmC analysis, Michael Ortiz and the CRI Flow cytometry facility for flow cytometry support, Dieu Linh Nguyen, Landon Nguyen, and Qing Ding for mouse colony management and technical assistance, Anjana Rao and Ageliki Tsagaratou for sharing *Tet3^fl^* mice, and Cheng Cheng Zhang for sharing MV-4-11 cells.

---

**Supplemental Figure 1.**
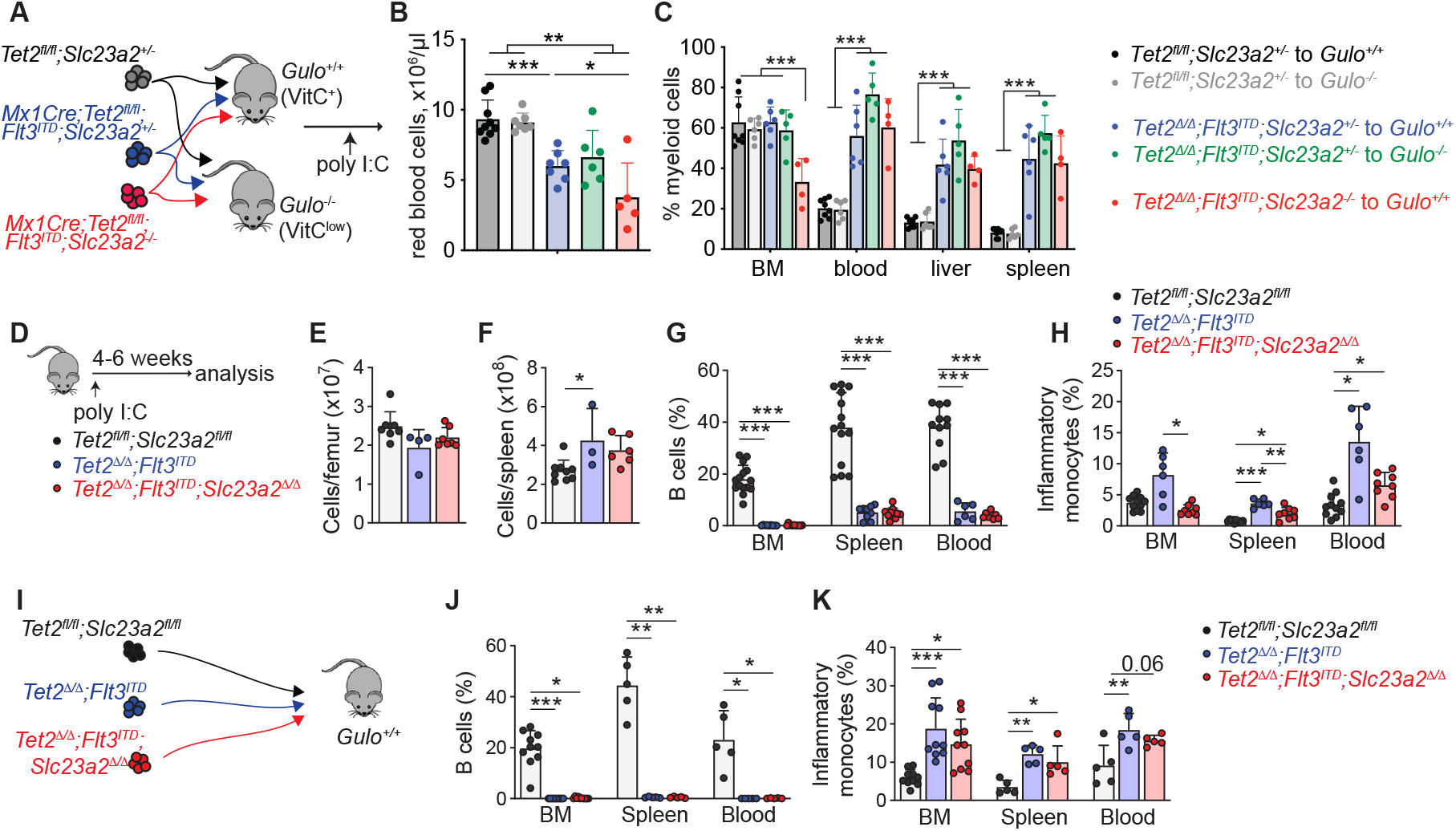
Ascorbate depletion promotes leukemogenesis. (**A-C**) Reciprocal transplantation of ascorbate-replete or ascorbate-depleted leukemia cells into ascorbate-replete or ascorbate-depleted mice and hematopoietic analysis. (**D-H**) Hematopoietic analysis of mice from the indicated genotypes. (**I-K**) Hematopoietic analysis of mice transplanted with ascorbate-replete or ascorbate-depleted leukemia cells. Statistical significance was assessed with a one-way ANOVA (B-C, F and K), Kruskal-Wallis test (E, G, BM; and J, BM), or a Brown-Forsythe ANOVA (G, spleen and blood; H and J, spleen and blood).

**Supplemental Figure 2.**
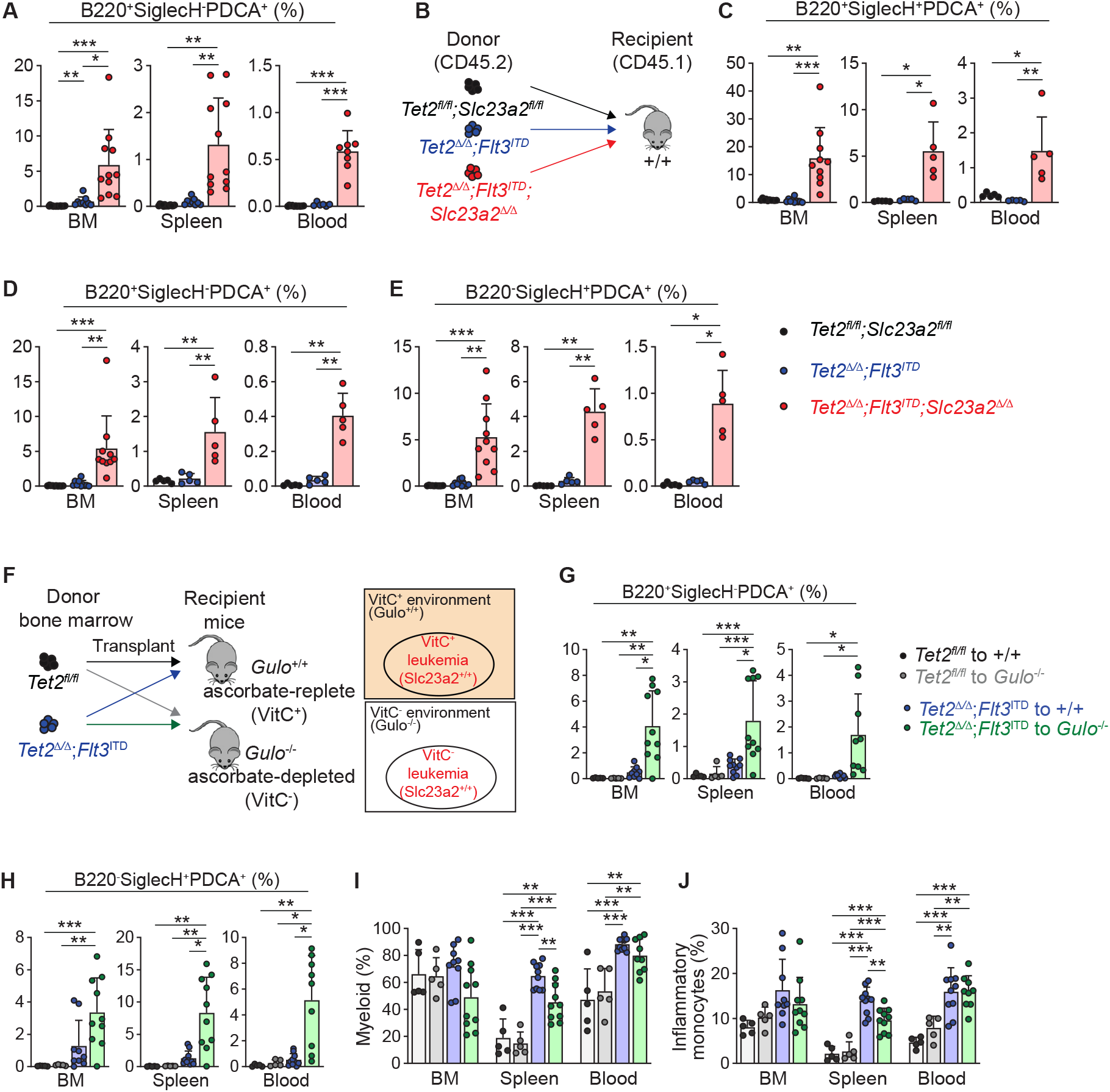
Ascorbate depletion promotes the development of plasmacytoid dendritic cells. (**A-E**) Frequencies of pDC populations in primary mice and in mice transplanted with ascorbate-replete or ascorbate-depleted leukemia cells. (**F-J**) Transplantation of leukemia cells into ascorbate-depleted or ascorbate-replete *Gulo^−/−^* mice followed by analysis 5 weeks later. Statistical significance was assessed with a Kruskal-Wallis test (A, BM; C, BM; D, BM; E, BM; G, spleen; and H, BM), Brown-Forsythe ANOVA (A, spleen and blood; C, spleen; D, blood; E, spleen and blood; G, BM and blood; and H, spleen and blood), or one-way ANOVA (C, blood; D, spleen; and I-J).

**Supplemental Figure 3.**
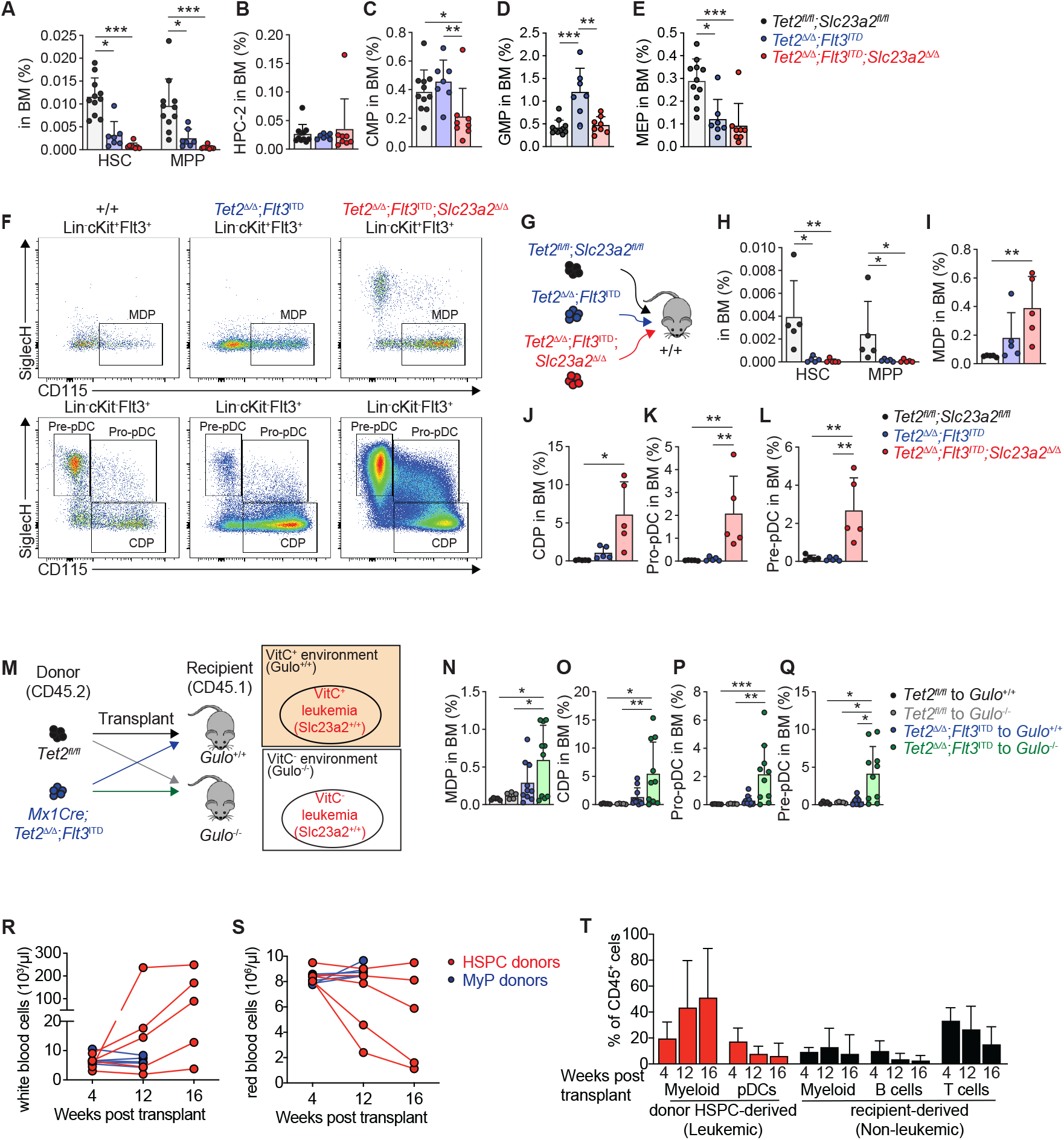
Analysis of leukemic stem and progenitor cells in ascorbate-depleted AML. (**A-E**) Analysis of HSPCs in primary mice. (**F**) Representative flow plots of pDC lineage progenitors in the bone marrow. (**G-L**) Analysis of HSPCs in transplant recipient mice. (**M-Q**) Frequencies of pDC lineage progenitors 5 weeks after transplant of *Tet2^Δ/Δ^*;*Flt3^ITD^* or control bone marrow cells into irradiated *Gulo^+/+^* or *Gulo^−/−^* mice. (**R-S**) White blood cell and red blood cell counts in individual transplant recipients. (**T**) The frequency of donor-or recipient-derived cell populations in the blood (n = 4 mice). Statistical significance was assessed with a Kruskal-Wallis test (A, C-E and H, HSC; and O-P), one-way ANOVA (H, MPP; I and K-L), or a Brown-Forsythe ANOVA (J, N and Q).

**Supplemental Figure 4.**
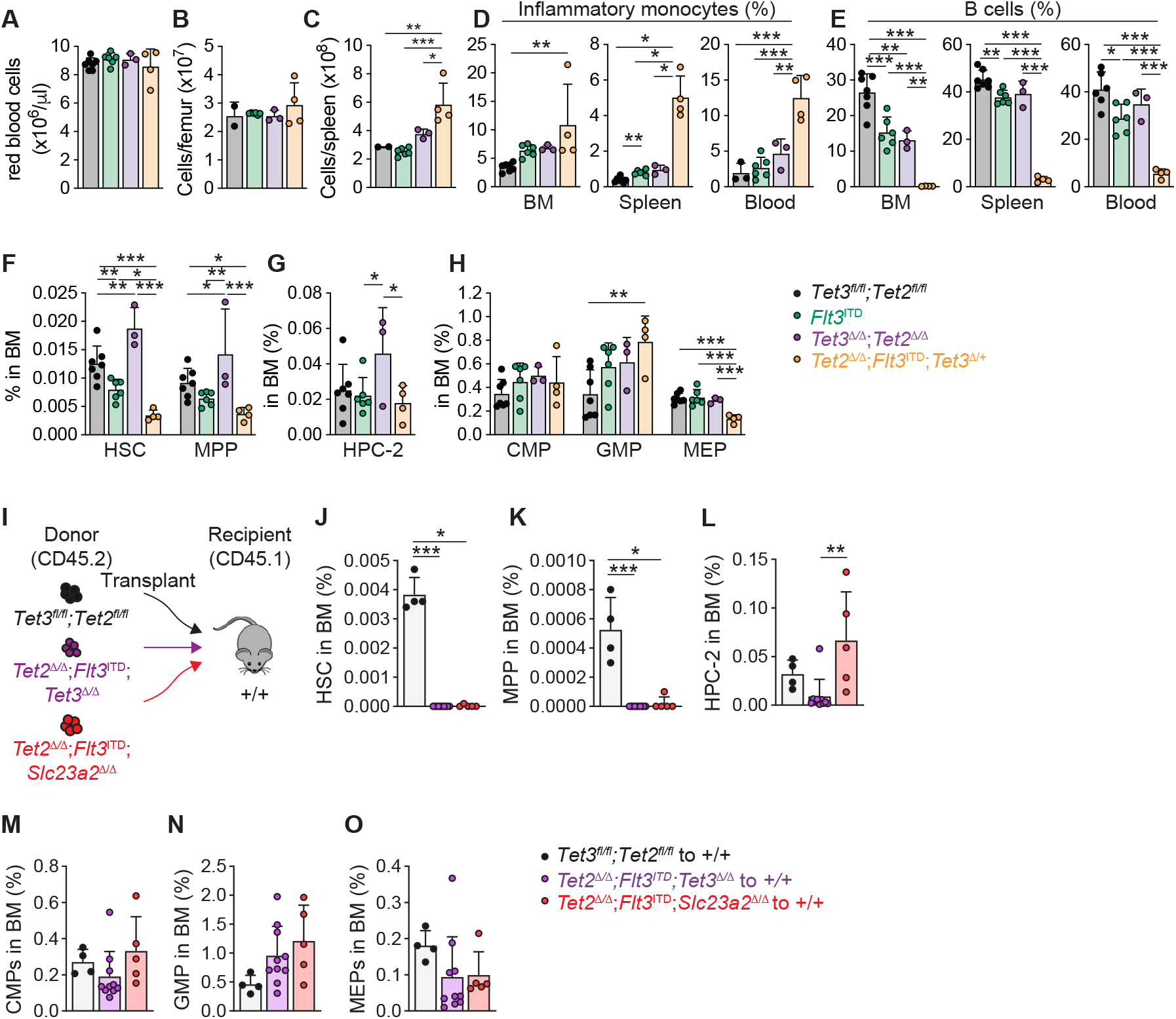
Ascorbate suppresses leukemia development through Tet2 and Tet3. (**A-H**) Hematopoietic analysis of *Tet2^Δ/Δ^;Tet3^Δ/+^;Flt3^ITD^*mice and littermate controls 4-6 weeks after deletion including: (**A-E**) analysis of mature cells; (**F-H**) analysis of HSCs and progenitors. (**I-O**) Analysis of HSCs and progenitors in transplant recipients of *Tet2^Δ/Δ^*;*Flt3^ITD^*;*Tet3^Δ/Δ^*or *Tet2^Δ/Δ^*;*Flt3^ITD^*;*Slc23a2^Δ/Δ^* or wild-type control bone marrow. Statistical significance was assessed with a one-way ANOVA (C, D, BM; E-H and N), Brown-Forsythe ANOVA (D, spleen and blood), or a Kruskal-Wallis test (J-M and O).

**Supplemental Figure 5.**
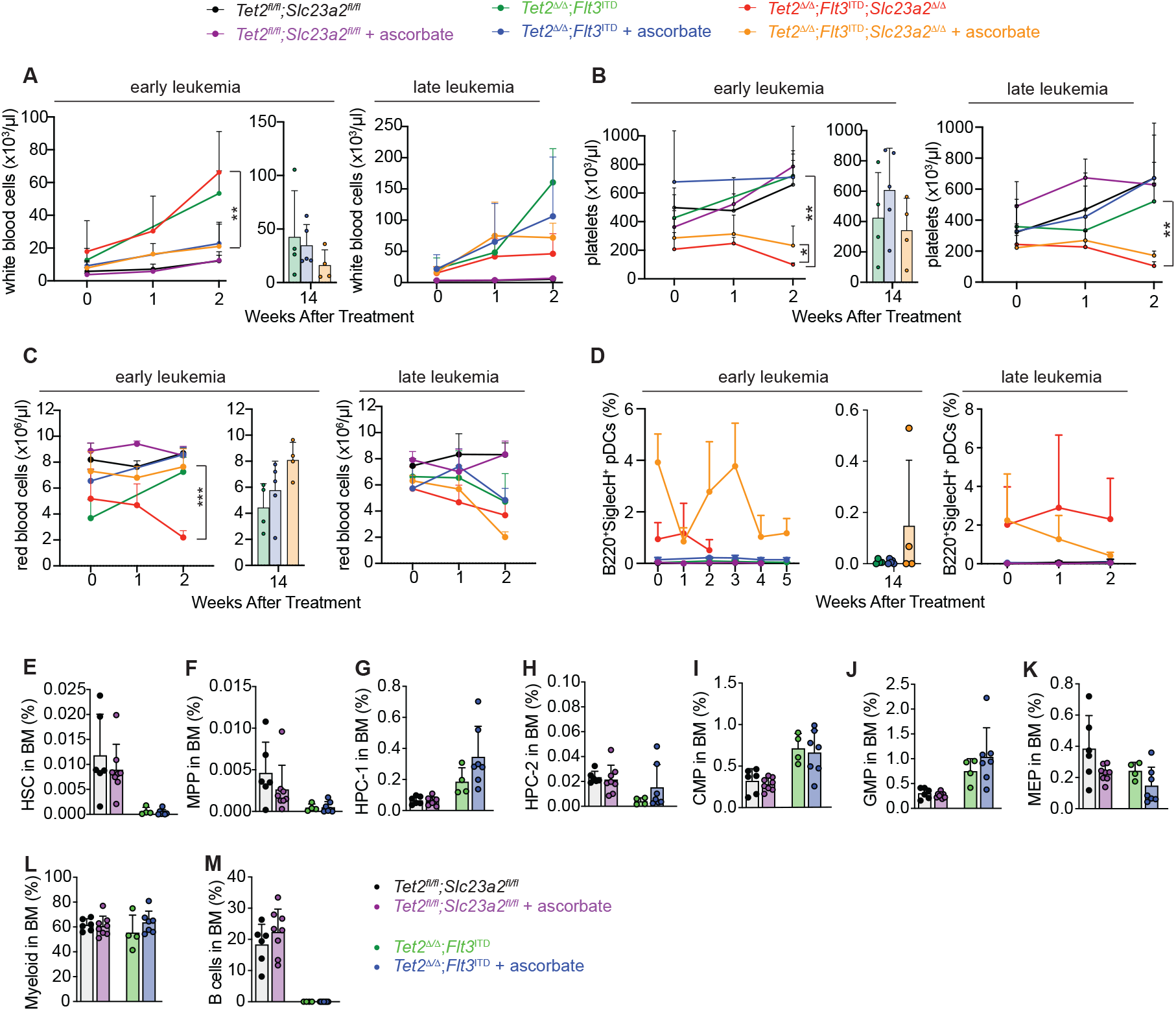
Effects of ascorbate administration on hematologic parameters in leukemia or on normal and *Tet2^Δ/Δ^*;*Flt3^ITD^* hematopoiesis. (**A-D**) Blood analysis of early-state or late-stage leukemia treated with ascorbate. (E-M) Hematopoietic analysis of mice transplanted with *Tet2^fl/fl^;Slc23a2^fl/fl^*, or *Tet2^Δ/Δ^*;*Flt3^ITD^* marrow cells and treated with ascorbate for 6-8 weeks. Statistical significance was assessed with a t-test (A and C), or Welch’s t-test (B).

**Supplemental Figure 6.**
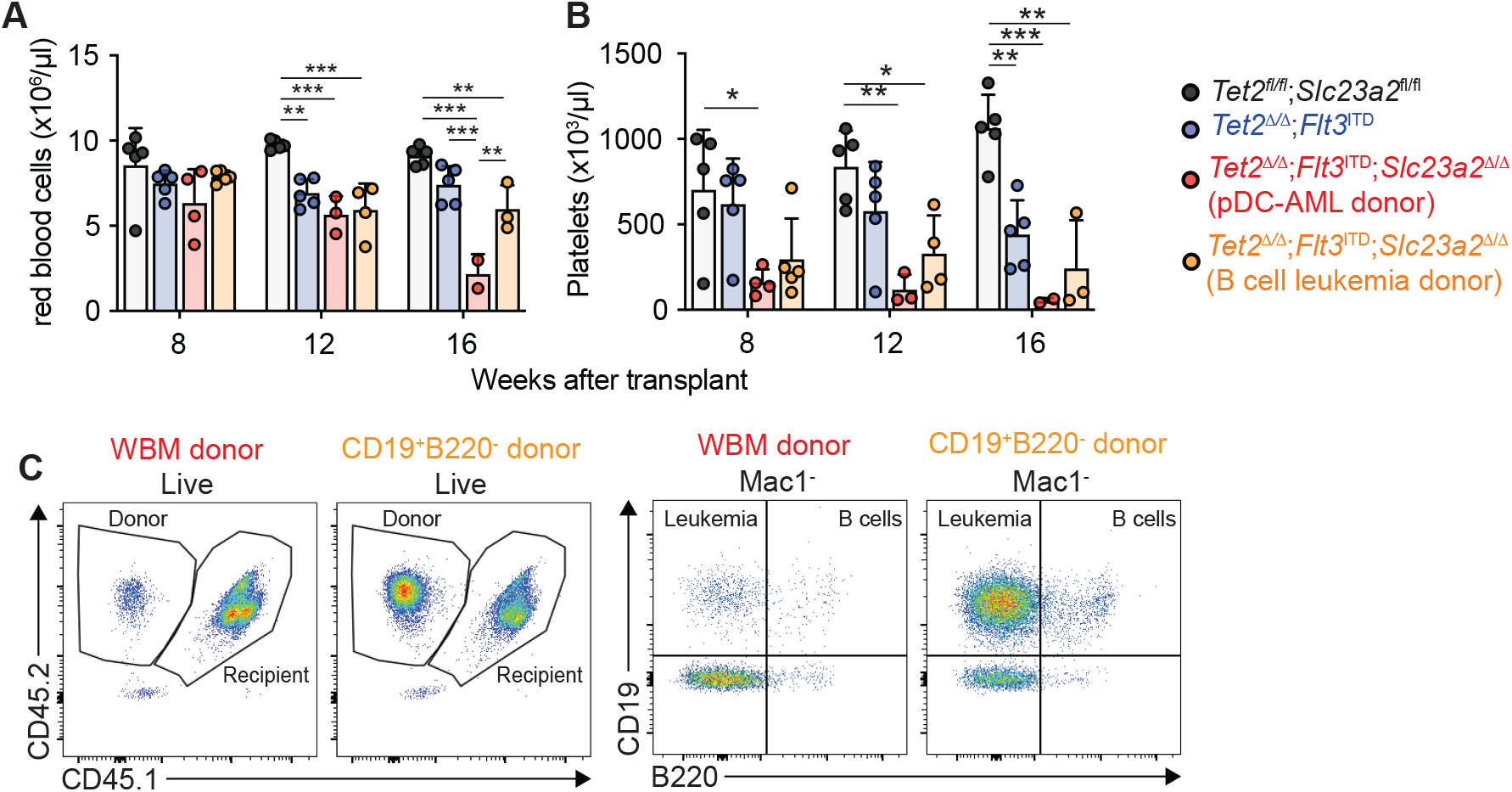
Analysis of *Tet2^Δ/Δ^;Flt3^ITD^;Slc23a2^Δ/Δ^* pDC-AML or B leukemia. (**A-B**) Red blood cell and platelet counts in recipient mice transplanted with bone marrow from mice with *Tet2^Δ/Δ^*;*Flt3^ITD^*;*Slc23a2^Δ/Δ^*pDC-AML or B cell leukemia. (**C**) Representative flow plots of leukemic CD19^+^B220^−^ cells in secondary transplant recipient mice after transplant with whole bone marrow cells or sorted CD19^+^B220^−^ cells from primary recipients of *Tet2^Δ/Δ^;Flt3^ITD^;Slc23a2^Δ/Δ^* cells. Statistical significance was assessed with a one-way ANOVA (A-B).

